# Pollinator and host sharing lead to hybridization and introgression in Panamanian free-standing figs, but not in their pollinator wasps

**DOI:** 10.1101/2022.08.04.502839

**Authors:** Jordan D. Satler, Edward Allen Herre, Tracy A. Heath, Carlos A. Machado, Adalberto Gómez Zúñiga, K. Charlotte Jandér, Deren A. R. Eaton, John D. Nason

## Abstract

Obligate pollination mutualisms, in which plant and pollinator lineages depend on each other for reproduction, often exhibit high levels of species-specificity. However, cases in which two or more pollinator species share a single host species (host sharing), or two or more host species share a single pollinator species (pollinator sharing), are known to occur in current ecological time. Further, evidence for host switching in evolutionary time is increasingly being recognized in these systems. The degree to which departures from strict specificity differentially affect the potential for hybridization and introgression in the associated host or pollinator is unclear. We addressed this question using genome-wide sequence data from five sympatric Panamanian free-standing fig species (*Ficus* subgenus *Pharmacosycea*, section *Pharmacosycea*) and their six associated fig pollinator wasp species (*Tetrapus*). Two of the five fig species, *F. glabrata* and *F. maxima*, were found to regularly share pollinators. In these species, ongoing hybridization was demonstrated by the detection of several first-generation (F1) hybrid individuals and historical introgression was indicated by phylogenetic network analysis. In contrast, although two of the pollinator species regularly share hosts, all six species were genetically distinct and deeply divergent, with no evidence for either hybridization or introgression. This pattern is consistent with results from other obligate pollination mutualisms, suggesting that, in contrast to their host plants, pollinators appear to be reproductively isolated, even when different species of pollinators mate in shared hosts.

## Introduction

Hybridization and introgression have contributed to the evolution of species and clades across the tree of life (Anderson, 1953; Stebbins, 1959; Mallet, 2005; Mallet et al., 2016). Genome-scale data and advances of analytical techniques, in particular, have enabled the detection and documentation of hybridization and introgression in many groups, and suggest that these processes are more widespread than previously thought (Taylor and Larson, 2019). These processes can be important contributors to speciation and adaptive radiations, which can be spurred by the introduction of beneficial alleles and multilocus combinations to a recipient lineage (adaptive introgression; *e.g*., Hedrick, 2013; Edelman and Mallet, 2021). Adaptive introgression has been documented in a diversity of lineages, including cichlid fishes (Meier et al., 2017; Malinsky et al., 2018; Svardal et al., 2020), butterflies (Pardo-Diaz et al., 2012; Enciso-Romero et al., 2017; Edelman et al., 2019), and oaks (Eaton et al., 2015; McVay et al., 2017; Leroy et al., 2020). Interspecific gene flow can also contribute to adaptive responses to rapidly changing environments, such as human-caused environmental modifications (Hamilton and Miller, 2016). Consequently, there is growing appreciation for the importance of hybridization and introgression in generating and maintaining organismal adaptation and diversity.

Flowering plants and their animal pollinators provide useful case studies for showing how ecological interactions can affect gene flow patterns and evolutionary trajectories (*e.g*., reproductive isolation or introgression in the associated host and pollinator lineages). Of particular interest, brood pollination mutualisms consist of host and pollinator lineages that obligately depend on each other for reproduction and survival. Examples of these highly specialized interactions include figs and fig wasps, yuccas and yucca moths, leaffiowers and leaffiower moths, globeflowers and globeflower flies, and palms and weevils (Cruaud et al., 2012; Hembry and Althoff, 2016; de Medeiros and Farrell, 2020; Pellmyr et al., 2020). Brood pollination mutualisms often exhibit strict host specificity (*i.e*., only one pollinator species is consistently associated with only one host species), which is thought to promote reproductive isolation for both the host and the pollinator.

Several studies, however, have revealed that two or more pollinator species per host (host sharing), or two or more host species per pollinator (pollinator sharing), are not uncommon in some systems (Molbo et al., 2003; Cornille et al., 2012; McLeish and Van Noort, 2012; Starr et al., 2013; Yang et al., 2015; Wang et al., 2016). These reduced specificity ecological interactions, along with accumulating molecular evidence of historical host switching (*e.g*., Cruaud et al., 2012; Hembry et al., 2013; Satler et al., 2019), are consistent with opportunities for hybridization in either the host, the pollinator, or both (Berg, 1989; Leebens-Mack et al., 1998; Machado et al., 2005; Cornille et al., 2012; Rentsch and Leebens-Mack, 2012; Wang et al., 2016; Arteaga et al., 2020; Wang et al., 2021a). This raises the more general question of how different patterns of host specificity affect the evolutionary dynamics (*e.g*., hybrid formation and introgression versus reproductive isolation) for each partner species in obligate plant–pollinator mutualisms. These dynamics, in turn, will affect the processes of speciation and diversification in both taxa.

Figs (*Ficus*, family Moraceae) and their pollinator wasps (family Agaonidae) represent an ancient (*≥*80 Ma) and diverse (*≥*900 described species of figs) obligate pollination mutualism (Janzen, 1979; Machado et al., 2001; Weiblen, 2002; Cook and Rasplus, 2003; Cruaud et al., 2012; Wang et al., 2021a). Pollination in this keystone mutualism results in fruit production that supports diverse frugivores across tropical and subtropical habitats worldwide (Shanahan et al., 2001). When receptive, the enclosed fig inflorescences (syconia) produce volatile chemicals that attract mated pollen-bearing female fig wasps (Van Noort et al., 1989; Ware et al., 1993; Grison-Pigé et al., 2002; Hossaert-McKey et al., 2010; Cornille et al., 2012; Wang et al., 2021b). These foundress wasps enter the syconia to pollinate flowers and oviposit in a subset of them. The pollinated flowers usually develop as viable seeds but those flowers that receive wasp eggs usually become galls that support the development of the wasp offspring. After maturing, the pollinator wasp offspring then mate within their natal fig syconia before females collect pollen and disperse—typically several kilometers (Herre, 1989; Nason et al., 1998; Ahmed et al., 2009)—to locate receptive syconia on other fig trees.

Fig pollinating wasp species often exhibit high levels of specificity to host fig species (Weiblen, 2002; Herre et al., 2008), which has contributed to the paradigm of one wasp to one fig. However, with increasingly deep spatial and temporal sampling, and the application of more sophisticated genetic techniques, it is increasingly clear that there is often considerable deviation from this paradigm. For example, several studies (based primarily on COI mtDNA barcode data) suggest some level of pollinator sharing or host sharing, and also imply host switching (*e.g*., Molbo et al., 2003; Machado et al., 2005; Cornille et al., 2012; Darwell et al., 2014; Yang et al., 2015; Wang et al., 2016; Yu et al., 2019). Further, roughly 30% of fig species have been estimated to host multiple pollinator species at either local or regional scales (Yang et al., 2015). This contemporary host sharing is complemented by detailed studies documenting an evolutionary history of host switching (Satler et al., 2019). These departures from strict one wasp species to one fig species interactions can potentially introduce heterospecific pollen to non-natal host fig species, allow individuals of heterospecific wasps to develop and potentially mate within the same individual fig syconium, or both. These observations all motivate the question—to what degree are hybridization and introgression observed in host figs or pollinator wasps?

There is morphological and molecular evidence for the existence of successful natural hybridization and introgression among closely related fig species within specific fig sections (*e.g*., Berg, 1989; Machado et al., 2005; Jackson et al., 2008; Wilde et al., 2020). This appears to be the case even among distantly related fig species across distinct subgenera (Compton, 1990; Ramírez, 1994; Wang et al., 2021a). In contrast to their fig hosts, however, the few studies conducted to date using microsatellites or even deeper genomic tools find little or no evidence of hybridization or successful introgression between co-occurring pollinator wasp species (Molbo et al., 2003, 2004; Sutton et al., 2017; Satler et al., 2022). Importantly, no study has directly applied genomic tools that can reveal the presence of hybridization and introgression across both the host figs and their associated pollinator species comprising an entire local community. Therefore, it is unclear the degree to which pollinator sharing, host sharing, host switching, or some combination of the three, generate hybridization and introgression in the host figs or pollinator wasps, and what these different ecological interactions mean for the strengthening or weakening of species boundaries.

Here we use genome-wide sequence data to test for hybridization and introgression in all species comprising a community of Panamanian free-standing fig hosts (five species of *Ficus* subgenus *Pharmacosycea*, section *Pharmacosycea*) and their associated pollinating fig wasps (six species of *Tetrapus*). We ask if hybridization and introgression have been operating within either the host figs, pollinator wasps, or both, and if so, whether currently observed levels of species specificity (or lack there of) explain these evolutionary processes. For the pollinating wasps, we find no evidence for hybridization or introgression among any of the *Tetrapus* species. We do identify, however, at least two pollinator species that are consistently pollinating and reproducing in more than one host fig species. For the host figs, we find that host species that frequently share the same pollinator species exhibit evidence of recent hybridization events (genetically identified F1 hybrids that are morphologically intermediate between parental types), as well as of historical introgression between two host fig species that are currently generating F1 hybrids. We discuss the implications of these findings for evolutionary dynamics of speciation in the fig hosts and wasp pollinators and discuss how the processes shaping fig and fig wasp evolution are consistent with observations from other obligate pollination mutualisms.

## Materials and Methods

### Wasp sampling and sequencing

We sampled pollinator wasps from the free-standing fig community located in central Panama in the vicinity of the Barro Colorado Island Nature Monument (BCNM, Table S1). The fig species that comprise this community are *F. glabrata, F. insipida, F. maxima, F. tonduzii*, and *F. yoponensis*. These five morphologically distinct species are native to our central Panama study area (Croat, 1978) and comprise roughly one-quarter of the 22 described species of Neotropical free-standing figs (Berg, 2006). Between March 2015 and February 2019, pollinator wasps were sampled from these five host fig species for genome-wide sequence analysis.

Mature fig syconia were brought to the lab on Barro Colorado Island where wasps were allowed to emerge in vials. To ensure independence among samples, one pollinator from each fig syconia was sampled and preserved in 95% EtOH or RNALater. Additionally, two wasp samples were also collected from sticky traps located near receptive figs of *F. tonduzii*. In total, we sampled 57 individual wasps representing six fig pollinator species from this community.

DNA was extracted using a Qiagen DNeasy blood and tissue kit (Qiagen Inc., Valencia CA, USA). Illumina libraries were generated using a KAPA Hyper prep kit with custom indices as described in Glenn et al. (2019). Samples were sheared using a Covaris sonicator to an average size of 400–500 base pairs. Following library prep, we grouped samples into sets of eight and conducted probe hybridization targeting 2590 ultraconserved element (UCE) loci using the hymenopteran probe v2 set of Branstetter et al. (2017). Size distributions were assessed with a Bioanalyzer and samples were grouped in equimolar concentrations for sequencing. We sequenced libraries on an Illumina sequencer (HiSeq 3000 and HiSeq 4000) generating 150 bp paired-end reads.

DNA sequence reads were processed with Phyluce v1.6.7 (Faircloth, 2015). Raw sequence reads were first processed with illumiprocessor v2.0.9 (Faircloth, 2013), a tool that uses Trimmomatic v0.39 (Bolger et al., 2014), to remove adapter contamination and poorly sequenced base pairs. Trinity v2.0.6 (Grabherr et al., 2011) was used to assemble cleaned reads into contigs. We then aligned contigs with the hymenopteran probe set v2 to retain only sequences matching a targeted UCE. Loci were aligned with MAFFT v7.407 (Katoh and Standley, 2013) and ends with high amounts of missing data were trimmed. Ambiguously aligned sites were removed with Gblocks v0.91b (Castresana, 2000) using default settings. We then filtered the cleaned sequence loci to retain those sampled in a minimum of 70% of individuals. We also generated a phased data set for the UCE loci following the outline of Andermann et al. (2018). Briefly, we aligned our cleaned sequence reads back to aligned loci with BWAMEM as implemented in bwa v0.7.17 (Li and Durbin, 2010). Data were phased using the phase command in samtools v1.9 (Li et al., 2009) resulting in two alleles per individual per locus. Phased data sets were cleaned as outlined above, and loci with a minimum of 70% of individuals were once again saved for downstream analysis.

### Wasp population structure and hybridization

We used two approaches to test for species boundaries and hybridization in the fig wasps. First, we used principal components analysis (PCA) to determine species groupings. In the absence of recent hybridization and introgression, individuals are expected to form distinct clusters corresponding to species. Hybrid individuals (F1s or subsequent backcrosses), in contrast, are expected to be located equidistant between species clusters while limited introgression is expected to result in intermixed species clusters. We conducted the PCA in R v3.6.3 (R Core Team, 2018) using the dudi.pca command in adegenet v2.1.3 (Jombart, 2008). For our input data set, we subsampled a single biallelic SNP per UCE locus. Missing data were replaced with the global mean allele frequency for that SNP. Because of a high amount of missing data for one individual (FW514), we excluded this sample from the analysis. We visualized results by plotting the first two principal component axes of variation.

Second, we explicitly tested for hybridization and introgression using the population graph and admixture approach as implemented in TreeMix v1.13 (Pickrell and Pritchard, 2012). TreeMix estimates a population graph with an *a priori* number of migration events between lineages, here species. This method allowed us to test if a model with migration between wasp species is a better fit to the data than a strictly bifurcating model without migration. We analyzed the data in TreeMix using zero to three interspecific migration events, and used the proportion of variance explained by the model to determine the optimal number of migration events. We used the phased data set and randomly subsampled a single biallelic SNP per locus for this analysis. If hybridization and introgression are not operating within this system, we would expect negligible improvement to the model as we add migration events.

### Wasp phylogenetics

To infer wasp phylogenetic relationships, we estimated a maximum likelihood (ML) phylogeny of the concatenated UCE data set in IQ-TREE v2.1.2 (Nguyen et al., 2015; Chernomor et al., 2016). This approach allowed us to determine if individuals sampled from the same fig host species cluster together in phylogenetic space. We partitioned the concatenated data set by UCE locus and used ModelFinder (Kalyaanamoorthy et al., 2017) with Bayesian Information Criteria (BIC) to select the substitution model of best fit for each partition. We assessed nodal support by generating 1,000 bootstrap replicates with the ultrafast bootstrap approximation (Hoang et al., 2018).

### Wasp mitochondrial DNA

We wanted to compare phylogenetic patterns between the nuclear (UCE) and mitochondrial genomes. If individuals belonged to different clades between the species tree (estimated with nuclear UCE data) and the mitochondrial gene tree, the cytonuclear discordance could be explained by interspecific hybridization and introgression. To generate mtDNA data from our samples, we followed the outline of (Satler et al., 2022). Briefly, we used NOVOPlasty v4.3.1 (Dierckxsens et al., 2017) to identify mitochondrial reads and generated haplotypes from off-target reads present in the UCE sequencing files. We used a COI sequence from a *Tetrapus* species (AY148155) as our seed sequence. After recovering mtDNA haplotypes, we aligned these data with MAFFT v7.471 and trimmed the matrix to match the length of the seed sequence to minimize missing data. We then estimated a ML gene tree with IQ-TREE, used ModelFinder with BIC to select the substitution model of best fit, and generated 1,000 bootstrap replicates with the ultrafast bootstrap approximation. Finally, we tested for cytonuclear discordance by comparing the species compositions recovered with the mitochondrial DNA with those recovered with the nuclear (UCE) DNA.

### Host associations

Through the estimation of well-supported wasp and host fig phylogenies, as well as knowledge of the fig species from which wasps were sampled, we can determine the association between pollinator and host fig species. A one-to-one correspondence between a wasp species and an individual host species is indicative of current host specificity. In contrast, evidence of reduced specificity is indicated when two or more host lineages share the same pollinators or when two or more wasp lineages are associated with the same host. This information can be quantified over available wasp samples to estimate the frequency with which each wasp species is associated with each fig species and to identify those wasp and fig species that have higher or lower host specialization. Lower host specificity creates greater opportunities for interspecific interactions between wasps and between figs, and provides a mechanism for hybridization.

Because our approach prioritized deep genomic sampling of individuals over sampling large numbers of individuals, we supplemented our data set with additional wasp individuals to increase sample size for assessing the host specificity of the different pollinator wasp species. Specifically, we collected COI mtDNA data from an additional 212 wasps sampled from the fig species described above (with the exception of *F. tonduzii*). Genomic DNA was extracted and COI sequences generated and aligned following the methods described in (Marussich and Machado, 2007). These COI mtDNA data provide sufficient information for confirming species identification and for generating host association frequencies.

This independent COI data set was generated from samples collected between February 1997 and May 2005, earlier than the samples collected here for UCE sequencing. Because of potential pollinator turnover, we needed to confirm the wasp species sampled for the newer UCE data set and the older COI data set were the same. To confirm wasp species identity between the two data sets, we combined the older COI data set of 212 wasps with our newer COI data set recovered from NOVOPlasty, resulting in a total of 268 sequences. We realigned these data with MAFFT, then estimated an ML gene tree with IQ-TREE (as outlined above) to test for continuity of wasp species over these two time periods. Since the two data sets resulted in congruent wasp species inference, we used this information from the combined COI gene tree to determine host–pollinator association frequencies.

### Fig sampling and sequencing

We sampled 30 fig trees representing all five free-standing fig species present in our Panamanian *Ficus* community and that were sampled for pollinating wasps (Table S2). This included five trees that were putatively identified as *F. glabrata* X *maxima* hybrids and one tree identified as a *F. insipida* X *maxima* hybrid. Our initial hybrid identifications were based on intermediate leaf morphology and growth form. Genomic DNA was extracted using a modified CTAB protocol (Doyle and Doyle, 1987). Extractions were sent to Floragenex Inc. (Eugene, OR, USA) for restriction-site associated DNA (RAD) library preparation. Single-end RAD libraries were generated with the PstI restriction enzyme following standard protocol (Baird et al., 2008). Libraries were sequenced on an Illumina HiSeq 3000 using 100 bp single-end sequencing.

DNA sequence reads were processed with ipyrad v0.9.62 (Eaton and Overcast, 2020). No mismatches were allowed in barcodes when demultiplexing samples, with strict filtering used for removing any adapter contamination. Up to five low-quality base calls were allowed in a read. We used a clustering threshold of 85% sequence similarity when assembling reads into loci within species. Within individuals, we allowed up to 5% Ns and 5% heterozygous sites per locus. Alleles were clustered across individuals using an 85% sequence similarity threshold. For clustered loci, we allowed up to 20% SNPs, up to 20% heterozygous sites, and up to eight total indels. Data sets were output varying the amount of missing data depending on downstream application.

### Fig population structure and hybridization

We conducted a PCA to determine if fig species cluster in multivariate space and to identify any potential hybrid individuals located between species clusters. To reduce potential negative effects of missing data, we output loci sampled from at least 90% of individuals, and from these data, selected a single SNP per locus. We conducted the PCA in R as described above for the wasps, and visualized the first two axes of variation.

Based on the PCA of SNP data (see Results), we identified six individual fig trees as putative recent hybrids between *F. glabrata* and *F. maxima*, and identified one individual as a putative hybrid between *F. maxima* and *F. yoponensis*. To further evaluate hybridization, we used fastSTRUCTURE (Raj et al., 2014) to estimate population membership between pure species and potential hybrids. fastSTRUCTURE uses a variational Bayesian framework to approximate the *structure* (Pritchard et al., 2000) model for estimating population membership. We created two data sets for fastSTRUCTURE, one that included individuals of *F. glabrata, F. maxima*, and putative hybrids between the two species, and one that included individuals of *F. maxima, F. yoponensis*, and their putative hybrid. We ran each data set under a two-population model (*K* = 2). If individuals represent hybrids, we would expect them to show population membership in both clusters in their respective analysis. For fastSTRUCTURE, we used unlinked SNPs present in at least 50% of sampled individuals. To visualize results, we used the R package pophelper v2.3.1 (Francis, 2017).

Of the hybrid figs identified in our community, we next wanted to know if they were first-generation hybrids (F1) or first-generation backcrosses (BC1) to either parental species. Individuals backcrossing to a parental species are of interest because they provide a mechanism for the introgression of genetic material between species. To address this question, we used snapclust (Beugin et al., 2018) implemented in the R package adegenet. Snapclust is a maximum likelihood approach for assigning individuals to clusters, including both pure species and hybrids. Specifically, snapclust can model F1 hybrids as well as first- and second-generation backcrosses, allowing the identification of the specific generation for a sampled hybrid. We partitioned samples into the same two data sets as described for fastSTRUCTURE, and used the same genomic data as described for the PCA.

### Fig phylogenetics and introgression

To estimate the phylogenetic relationships among these free-standing figs, we used the coalescent-based approach SVDQuartets (Chifman and Kubatko, 2014) as implemented in PAUP* v4.0a168 (Swofford, 2003). SVDQuartets uses site patterns in the nucleotide data to estimate a phylogeny under the coalescent model. Because hybridization and introgression are not accounted for in this method, we removed hybrid individuals identified by the above analyses from the unlinked SNP data set. Individuals were assigned to species *a priori*, all quartets were evaluated, and nodal support values were generated with 100 standard bootstrap replicates. We included an individual from *F. obtusifolia* (Bioaccession #SAMN12175287), a species of Neotropical strangler fig (*Ficus* subgenus *Urostigma*, section *Americanae*), to serve as the outgroup to estimate the root position of the phylogeny.

Although SVDQuartets estimates phylogenetic relationships among species while accounting for incomplete lineage sorting (ILS), the method does not model gene flow (*i.e*., it explicitly considers distinct non-hybridizing species). To test if hybridization and subsequent introgression have been processes operating among these fig species at deeper time-scales, we used the maximum pseudolikelihood approach SNaQ (Solís-Lemus and Ané, 2016) as implemented in PhyloNetworks (Solís-Lemus et al., 2017). This approach estimates a multispecies network by modeling the processes of ILS and hybridization. Thus, we can test if a model allowing ILS and hybridization is a better fit to the data than a model only allowing ILS.

To estimate the phylogenetic network, we first generated concordance factors from our unlinked SNP data sampled from pure species (as in SVDQuartets, hybrids were removed) as outlined in Olave and Meyer (2020). Specifically, we used the R function SNPs2CF (www.github.com/melisaolave/SNPs2CF), sampled 100 alleles per species quartet (n.quartet=100), and generated 100 bootstrap replicates. Using these concordance factors as input to SNaQ, we estimated networks allowing a maximum (hmax) number of between zero and three hybrid edges (*i.e*., hybridization events), doing ten runs per analysis. For the network analysis with zero hybrid edges, we used the SVDQuartets species tree as the starting tree. For each subsequent analysis, we used the hmax - 1 network as our starting network. Pseudolikelihoods were compared across runs with different numbers of hybrid edges to estimate the network model with best support. We then used the best model to estimate 100 bootstrap replicates to generate support for the presence of the hybrid edge(s) indicating historical gene flow and introgression between species.

## Results

### Wasp population structure and hybridization

We generated 180,616,361 raw reads for the 57 fig wasp pollinator samples representing six pollinator species (Table S3). Individuals had on average 3,168,708 (*±* 1,603,597) raw reads. Following data processing, individuals had on average 154,359 (*±* 74,989) contigs with an average length of 379 (*±* 211) base pairs. We generated contig data from 2,248 total UCE loci, with each individual being sequenced at 1,423 (*±* 151) loci on average (Table S3).

All six wasp species are well differentiated with little intraspecific variation in PCA space (Figure 1). PC1 and PC2 explained 44.19% and 20.11% of the variation, respectively. Because we see tight clusters of individuals within species, and no spread of individuals between species, the PCA supports the pollinators as genetically distinct species with no recent hybridization or introgression.

**Figure 1:**
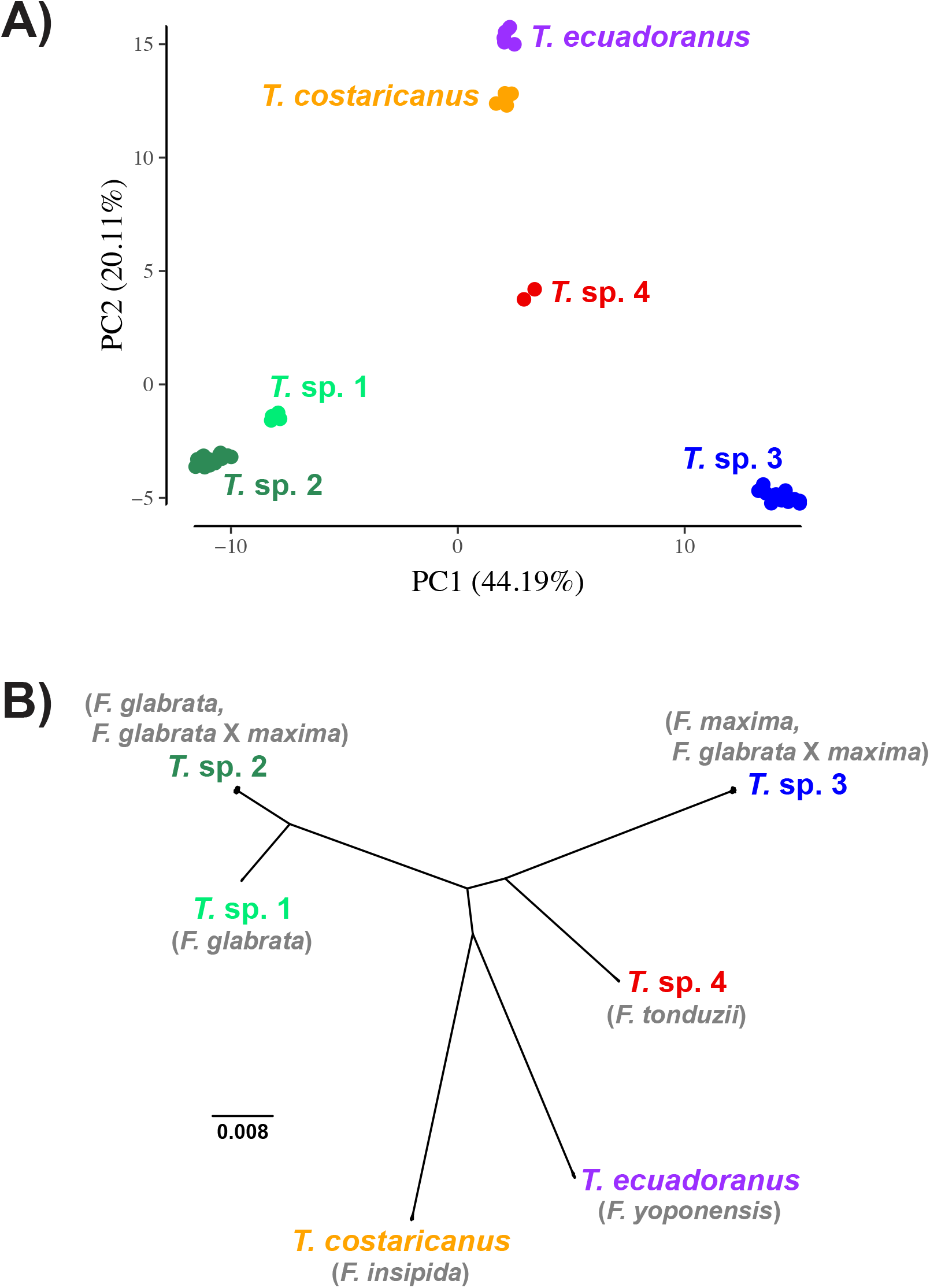
Population genetic and phylogenetic results for the fig pollinating wasps. A) Principal components analysis (PCA) of the *Tetrapus* wasps. Species are genetically distinct with little intraspecific divergence. B) An unrooted maximum likelihood phylogeny. All interspecific nodes are strongly supported with bootstrap values of 100. Fig host species names are shown in grey.

TreeMix results indicate that a model without hybridization provides the best fit to the data (Table 1). Although we estimated models with up to three admixture edges, there was essentially no increase in the proportion of variation explained by the model with the addition of admixture edges. Thus, the PCA and TreeMix results both show the pollinator species to be genetically distinct and reproductively isolated, with no evidence of hybridization.

**Table 1:** TreeMix results for the pollinator wasps. We compared models allowing between zero (m0) and three (m3) admixture edges. There is essentially no increase in the proportion of variation explained by the model when allowing admixture edges, suggesting a model with zero admixture edges is a best fit for the data. This is consistent with an absence of introgression among these pollinator wasp species.

| Model | Admixture Events | Percent Variation |
| --- | --- | --- |
| m0 | 0 | 99.87% |
| m1 | 1 | 99.99% |
| m2 | 2 | 99.99% |
| m3 | 3 | 99.99% |

### Wasp phylogenetics

In agreement with the PCA and TreeMix results, a concatenated ML tree of the UCE data detected six well-defined and reciprocally monophyletic species (Figure 1). The tree was characterized by low intraspecific divergence and high interspecific divergence. All interspecific nodes have 100 bootstrap support values.

### Wasp mitochondrial DNA

We were able to generate COI data from 56 of the 57 wasp individuals. After alignment and edge trimming, our COI matrix was composed of 816 base pairs. In agreement with the nuclear UCE phylogeny, our COI gene tree recovered six clades, all supported with bootstrap values of 100 (Figure S1). Once again, these species were characterized by low intraspecific divergence and high interspecific divergence. Results recovered with mtDNA data mirrored those recovered with UCE data, showing no cytonuclear discordance among species compositions between the two data sets.

### Host associations

Species compositions were identical between the UCE nuclear phylogeny and the mitochondrial gene tree. Pollinator species have also been consistent temporally in this community between the previously generated COI data and the current UCE data set. We therefore combined our current sampling with the 212 samples directly sequenced for COI to quantify associations between host figs and pollinator wasps in this community (Figure 2). Additionally, because we identified six individual figs as being recent hybrids between *F. glabrata* and *F. maxima* (see below), we grouped these individuals as a distinct host fig species (*F. glabrata* X *maxima*) for understanding host associations.

**Figure 2:**
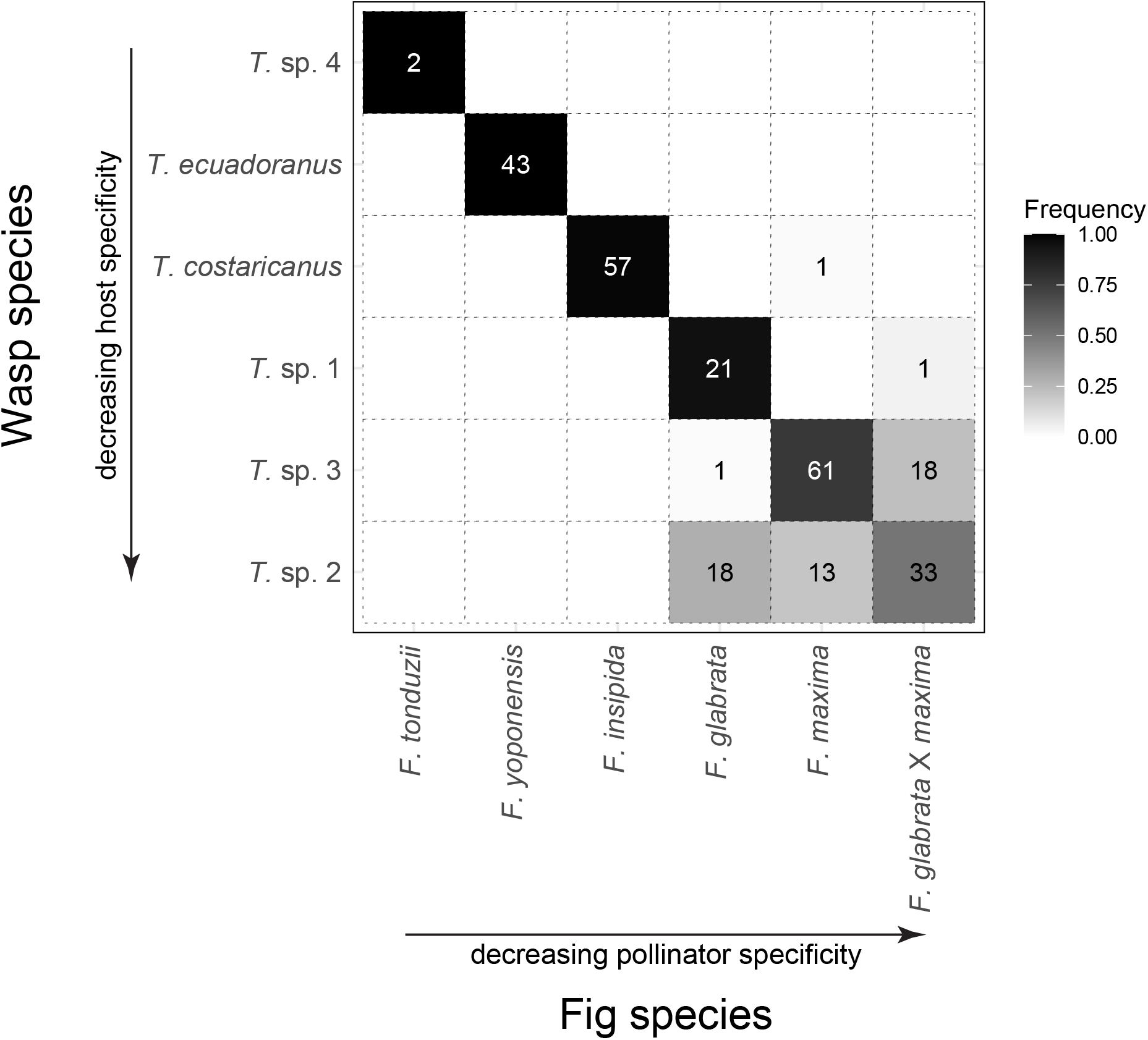
A heatmap showing the relative frequency with which each of the six *Tetrapus* wasp species was sampled from the five *Pharmacosycea* fig species for UCE and COI sequencing. Because we identified six individual figs as recent hybrids between *Ficus glabrata* and *F. maxima*, we treat them here as a distinct lineage for understanding host associations (see Results). Wasp and fig species are ordered from having higher to lower host or pollinator specificity, respectively. Rows with multiple entries represent cases of hosts sharing pollinators, which could promote hybridization between fig species. Columns with multiple entries represent cases of pollinators sharing hosts, which could promote wasp hybridization. Values in the cells are the numbers of wasps sampled per host combined over UCE and COI sequencing data sets.

Of the 269 wasps sampled, only three (1.1%) appear to be cases in which a single individual of a given wasp species pollinated a non-natal host species (one individual each of *T. costaricanus* on *F. maxima, T*. sp. 1 on *F. glabrata* X *maxima*, and *T*. sp. 3 on *F. glabrata*). Looking beyond these rare “mistakes” in host association, a single pollinator species is associated with each of *F. insipida* (*T. costaricanus*), *F. tonduzii* (*T*. sp. 4), and *F. yoponensis* (*T. ecuadoranus*), strongly suggesting strict host specificity in these species. In contrast, we also identified clear cases of host sharing, resulting in potential wasp species co-occurrence in the same figs of the same host species, and pollinator sharing, resulting in potential fig species being pollinated with interspecific pollen, and therefore opportunities for hybridization for both the figs and the wasps.

Specifically, two pollinator species are regularly associated with *F. glabrata*: one pollinator species is host specific (*T*. sp. 1), whereas the other species (*T*. sp. 2) is commonly associated with *F. glabrata, F. maxima*, and *F. glabrata* X *maxima* hybrids (Figure 2). Additionally, one pollinator species (*T*. sp. 3) is primarily associated with *F. maxima*, but individuals were also frequently sampled from *F. glabrata* X *maxima* hybrids. We thus have two cases of host sharing: (*T*. sp. 1) and (*T*. sp. 2) associated with *F. glabrata*, and (*T*. sp. 2) and (*T*. sp. 3) associated with *maxima* and also with *F. glabrata* X *maxima* hybrids (Figure 2). Moreover, we have two cases of pollinator sharing: *F. glabrata, F. maxima*, and *F. glabrata* X *maxima* hybrids share a pollinator (*T*. sp. 2), as do *F. maxima* and *F. glabrata* X *maxima* hybrids (*T*. sp. 3), which directly affects opportunities for hybridization among fig lineages. In sum, four of the pollinator species are host-species specific while two are associated with multiple hosts.

### Fig population structure and hybridization

We generated 263,931,402 raw reads for the 30 fig tree samples representing five fig species (Table S4). Individuals had on average 8,797,713 (*±* 7,271,731) raw reads. Following data processing, individuals had on average 76,879 (*±* 31,497) rad clusters (Table S4).

Requiring at least 90% coverage for loci, we used 9,662 unlinked SNPs for the PCA. PC1 and PC2 explained 25.09% and 22.29% of the variation, respectively. Species are recovered as distinct clusters in PCA space (Figure 3). We also recovered individuals that appear to represent genetic hybrids. For example, we recovered six individuals as a cluster (black squares) approximately equidistant between *F. glabrata* and *F. maxima*. In addition, another putative hybrid fig individual (grey triangle) is equidistant between *F. maxima* and *F. yoponensis*. The existence of multiple hybrids between *F. glabrata* and *F. maxima* is consistent with the pollinator sharing observed between these two species, and the intermediate positions of these admixed fig individuals are consistent with recent hybridization.

**Figure 3:**
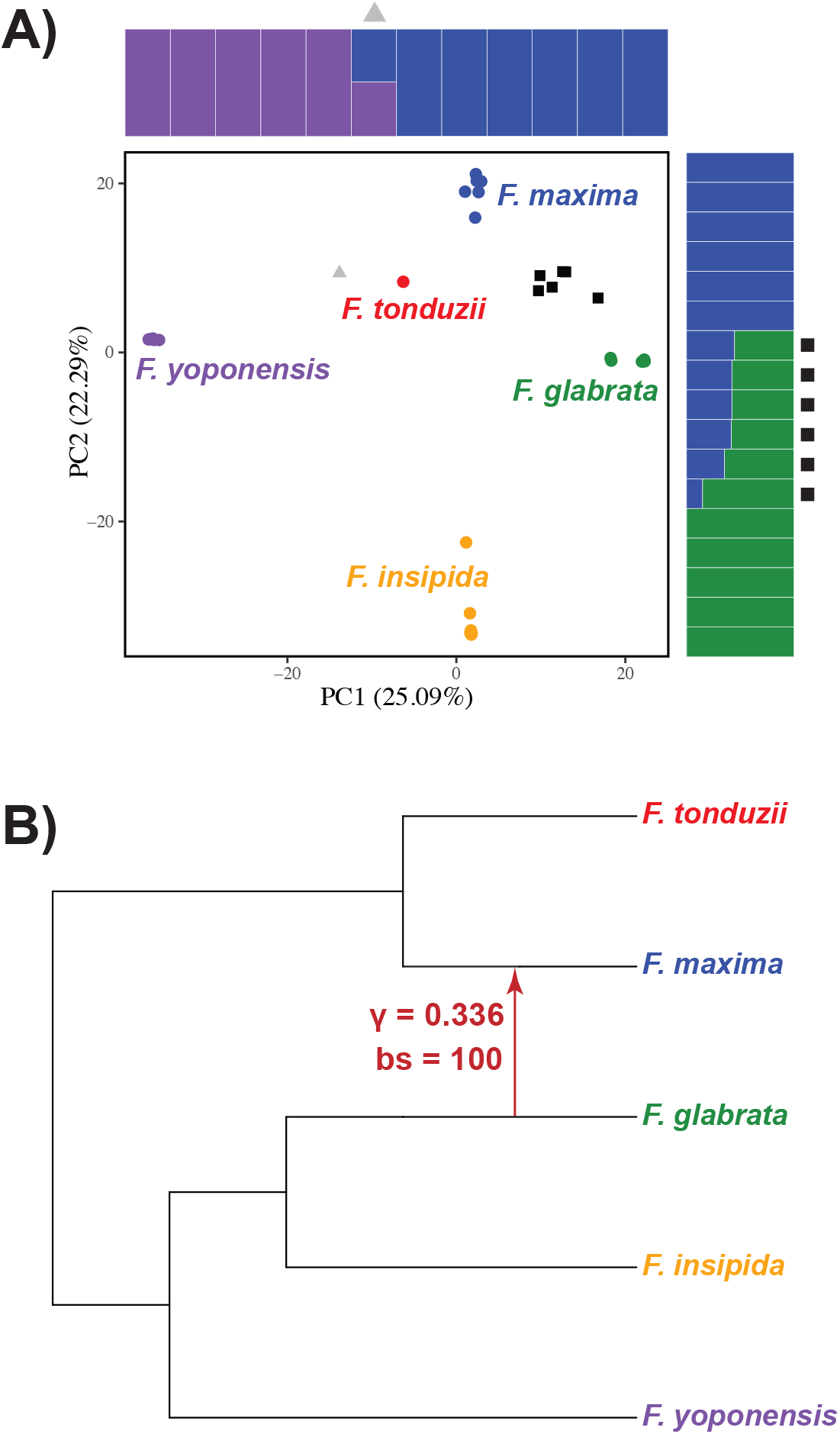
Population genetic and phylogenetic results for the host figs. A) Principal components analysis (PCA) of section *Pharmacosycea* figs. Squares in black represent hybrid individuals between *Ficus maxima* and *F. glabrata*, with the triangle in gray representing a hybrid individual between *F. maxima* and *F. yoponensis*. Genetic structuring plots from fastSTRUCTURE—color-coded to match corresponding species in the PCA—show hybridization between the species, with symbols (black squares and grey triangle) corresponding to individuals in the PCA plot. B) Phylogenetic network of the fig species. Hybrid individuals were removed from this analysis. The best model places one hybrid edge between *F. glabrata* and *F. maxima*, estimating that *F. maxima* inherited 33.6% of its genome from *F. glabrata*. All nodes and the hybrid edge are strongly supported with bootstrap values of 100. The outgroup (*F. obtusifolia*) was removed for visual purposes.

To further explore species affinities of the putative hybrid individuals, we analyzed subsets of the data in fastSTRUCTURE. One analysis contained samples from *F. glabrata, F. maxima*, and the putative hybrids of the two, while the other contained samples from *F. maxima, F. yoponensis*, and the putative hybrid between those two. For each analysis, the two pure species were recovered as distinct with the putative hybrid samples showing admixed ancestry consistent with recent hybridization (Figure 3). To estimate if these recent hybrids are first-generation hybrids (F1s) or first- or second-generation backcrosses, we analyzed the two data sets in snapclust. The result for the *F. glabrata* and *F. maxima* data set estimated that five individuals are F1 hybrids and one individual is a first-generation backcross (BC1) to *F. glabrata* (Figure S2). The results for the *F. maxima* and *F. yoponensis* data set estimated the putative hybrid individual to be an F1 hybrid (Figure S2). These results confirm the presence of seven recent hybrids (six F1s, one BC1) in our sampled fig community, with the remaining individuals assigned to pure species.

### Fig phylogenetics and introgression

Our species tree estimated with SVDQuartets recovers the fig species in two clades: one supporting *F. glabrata* and *F. insipida* as sister species, with *F. yoponensis* sister to them (all nodes strongly supported with a bootstrap value of 100), and a weakly supported sister relationship (bootstrap value of 51) between *F. maxima* and *F. tonduzii* (Figure S3). Although we sampled six hybrids between *F. glabrata* and *F. maxima*, demonstrating recent hybridization, these two species were not recovered as sister species in our phylogeny. This phylogenetic pattern is also the case with *F. maxima* and *F. yoponensis*, where they have produced an F1 hybrid but were not estimated to be sister species.

Because SVDQuartets does not explicitly consider hybridization and only models the process of ILS, we used a network approach to test if accounting for hybridization better reflects the evolutionary history of this community of fig species. Although we tested models with up to three hybrid edges, models with hmax > 1 always estimated a phylogenetic network with a single hybrid edge. While there was a drastic decrease in the pseudolikelihood from a model with zero hybrid edges (−3743.91) to a model with one hybrid edge (−1154.21), there was no decrease when allowing more hybrid edges. This suggests that a phylogenetic network with one hybrid edge best fit our data. This single hybrid edge was placed between *F. glabrata* and *F. maxima* with a bootstrap value of 100, and with 33.6% of the genome of *F. maxima* inferred to be inherited from *F. glabrata* (Figure 3). The phylogenetic relationships estimated in SNaQ are all strongly supported (bootstrap values of 100) and show the same pattern as relationships estimated with SVDQuartets.

## Discussion

We collected genome-wide sequence data from all species comprising a community of central Panamanian free-standing fig species and their associated pollinator wasp species. We used these data to assess evidence for hybridization and introgression in both the fig and pollinator taxa. Given the observed levels of pollinator and host sharing in this system, we applied rigorous genomic tests to determine whether associations with lower species specificity have contributed to hybridization and introgression in the host figs, the pollinator wasps, or both. For the host figs, we identified several individual F1 hybrids demonstrating hybridization at shallow time-scales, and recovered evidence for historical introgression at deeper time-scales between two species (*F. glabrata, F. maxima*) known to currently share pollinator species in central Panama. In contrast, we found no evidence of hybridization or introgression among any of the pollinator wasp species in either shallow or deep time-scales. These results suggest that pollinator sharing has generated genetic exchange between their host figs, blurring species boundaries. Yet, despite potential interspecific interactions resulting from sharing host species, reproductive isolation has apparently only been reinforced in the fig wasps.

### Pollinator sharing leads to hybridization in the figs

Pollinating fig wasps often appear to be highly species-specific in their associations with host figs (Ramírez, 1970; Bronstein, 1987; Moe et al., 2011; Satler et al., 2022). The degree to which this is true constrains the opportunities for hybridization (and subsequent introgression) in both lineages. Strict species-specificity by the pollinators necessarily limits potential interspecific pollination in their hosts. For host figs to have opportunities for hybridization, several barriers must be overcome by pollinators. A wasp bearing heterospecific pollen must disperse and recognize a species different from the one in which she developed (Compton, 1990; Nason et al., 1996). Once inside the syconium, the wasp must be able to put viable pollen grains in contact with the stigmatic surfaces of receptive flowers so that they can potentially produce viable F1 seeds.

However, even occasional or rare opportunities can be sufficient to promote genetic exchange between either the fig or pollinator species. For example, Moe and Weiblen (2012) tested for reproductive isolation among six sympatric dioecious fig species found in New Guinea. Using microsatellite data, they found 7 of 300 individual trees sampled to be of hybrid origin, demonstrating occasional hybrid formation even though pollinators are primarily host species-specific in the community (Moe et al., 2011). This shows that even limited opportunities for pollinator sharing and heterospecific pollen transfer can be sufficient to induce hybridization and, potentially, subsequent introgression in the host figs.

Of the six pollinator species sampled in the Panamanian community, two species are regularly associated with multiple fig species. One pollinator species (*T*. sp. 2) pollinates and successfully develops in *F. glabrata, F. maxima*, and *F. glabrata* X *maxima* hybrids, while a second (*T*. sp. 3) pollinates and successfully develops in *F. maxima* and *F. glabrata* X *maxima* hybrids. Pollinator sharing by these two wasp species predicts opportunities for hybrid formation between the host species, *F. glabrata* and *F. maxima*. This is exactly what we observe.

Of the thirty individual fig trees sampled, some appeared to be morphological intermediates, suggesting hybrid individuals. Our genetic tests confirmed this, identifying five F1s and one BC1 between *F. glabrata* and *F. maxima*. We then used a phylogenetic network approach restricting our data set to only individuals representing pure species. Using this method, the best model placed a hybrid edge indicating introgression between *F. glabrata* and *F. maxima* (Figure 3). These results demonstrate that hybridization and introgression between *F. glabrata* and *F. maxima* are both ongoing and have been operating at deeper time-scales, shaping the evolution of these two fig species. Although *F. glabrata* and *F. maxima* are genetically and morphologically distinct, incomplete reproductive isolation allows for hybrid compatibility, and thus, porous species boundaries in these two species. In addition, our genetic tests identified an F1 hybrid fig between *F. maxima* and *F. yoponensis*, and morphological intermediates between *F. insipida* and *F. yoponensis* have been observed, but unfortunately were not tested genetically (E. A. Herre, personal observation).

Evidence for hybridization and introgression have been detected in other sections of *Ficus*. In the Neotropics, Machado et al. (2005) and Jackson et al. (2008) used data from multiple loci to recover evidence supporting hybridization among several species of Panamanian strangler figs (*Ficus* subgenus *Urostigma*, section *Americanae*). Notably, a subset of these strangler figs in the Panamanian community are known to share pollinator species (Molbo et al., 2003; Machado et al., 2005; Jackson et al., 2008; Satler et al., 2019, 2022), providing a potential mechanism for interspecific pollination and hybridization. Sampling from a community of five dioecious fig species (*Ficus* subgenus *Sycomorus*, sections *Sycomorus* and *Hemicardia*) distributed in southeast Asia, Wang et al. (2016) found evidence of both pollinator sharing and host fig hybridization. In particular, they found 13.15% of sampled pollinator wasps were associated with non-natal fig host species, and 4.68% of fig individuals were of hybrid origin. In Australia, Wilde et al. (2020) found evidence of hybridization and subsequent backcrossing between two dioecious sandpaper fig species, *F. aculeata* and *F. coronulata*. Although the pollinator species is unknown for these two fig species, Wilde et al. (2020) suggests the fig hybrids provide another example of a breakdown of the one-to-one fig–pollinator association. Cytonuclear discordance also suggests historical introgression as an important process in the fig section *Galoglychia* (Renoult et al., 2009) and across species in even distantly related fig sections in general (Bruun-Lund et al., 2017). In particular, Wang et al. (2021a) analyzed whole-genome sequence data (including nuclear, mitochondrial, and chloroplast genomes) from lineages representing all major fig sections and recovered an extensive history of hybridization and introgression—both within and between sections—in *Ficus*. Although there are several prezygotic mechanisms potentially limiting fig hybridization, including the production of host-specific pollinator-attracting floral volatile blends (Van Noort et al., 1989; Ware et al., 1993; Grison-Pigé et al., 2002; Hossaert-McKey et al., 2010; Cornille et al., 2012; Wang et al., 2021b), these barriers appear to be less restrictive in the figs than in the pollinators. Our results are consistent with previous evidence and suggest that hybridization and introgression have been processes operating in shallow and deep time-scales in the evolution of the figs, but not in the evolution of their pollinator wasps. This asymmetry in relative importance of introgression appears to be an integral characteristic of the evolutionary history of the fig and fig wasp pollinator mutualism.

### Figs and their wasp pollinators differ in rates of successful hybridization

Evolutionary processes in the fig–pollinator mutualism appear to affect figs and wasps differently. While signatures of hybridization and introgression are present in host fig species (*e.g*., Machado et al., 2005; Renoult et al., 2009; Bruun-Lund et al., 2017; Wilde et al., 2020; Wang et al., 2021a), there is little evidence these processes affect fig wasp pollinators (see Molbo et al., 2003, 2004; Sutton et al., 2017; Satler et al., 2022). This suggests that the processes governing reproductive isolation operate differently within each lineage (figs– plants versus wasps–insects). Particularly when the one-to-one fig-to-wasp association is broken, many factors potentially contribute to the mechanisms allowing for hybridization and introgression within the figs, but apparently not within the pollinators.

As the pollinators of fig trees, fig wasps determine patterns of pollen gene flow between conspecific hosts, heterospecific hosts, or both. Whether a pollen-bearing wasp emerges from a given host species and then pollinates a different host species defines the opportunities for hybridization and introgression for their hosts. Thus, the ability of a pollinator—carrying heterospecific pollen—to detect, locate, enter, and successfully pollinate a non-natal fig species is critical for either reinforcing or blurring species boundaries. And because exceptions to the one-to-one fig-to-pollinator association are becoming more evident with increased taxon and genetic sampling (Darwell et al., 2014; Yang et al., 2015; Sutton et al., 2017; Souto-Vilarós et al., 2019; Yu et al., 2019), coupled with a growing appreciation of the role of host switching in these systems (Satler et al., 2019), it is probable that hybridization and introgression in the host figs is more widespread than previously realized. Indeed, an increase in genomic data sets has led to an increase in the detection of hybridization events across the tree of life (Taylor and Larson, 2019). We suggest that as more systems are explored with genome-scale data and are explicitly tested for hybridization and introgression, this will also be the case for figs (Wang et al., 2021a).

In our community of *Tetrapus* wasps, species are genetically distinct and are highly divergent (Figure 1). This result is consistent with genetic studies of fig wasps, where species typically show little intraspecific divergence but are deeply divergent from other species (*e.g*., Satler et al., 2022). The lack of hybridization in *Tetrapus* wasps is congruent with results from a recent study of Neotropical strangler fig pollinators. For example, Satler et al. (2022) sampled over 1,000 genome-wide UCE loci from a central Panamanian community of 19 *Pegoscapus* species associated with 16 strangler fig species. They recovered no signal of hybridization or introgression among these pollinators, even among pollinators known to share the same host fig species and to mate within the same fig syconia. In Australia, Sutton et al. (2017) sampled multiple pollinator species associated with the host fig *F. rubiginosa*. Even though 13% of figs had syconia with multiple pollinator species, they recovered no evidence of hybridization between pollinator species. Between results presented here and others (Sutton et al., 2017; Satler et al., 2022), studies that have explicitly tested for interspecific hybridization and introgression among fig pollinating wasps have yet to find evidence for these evolutionary processes. Thus, on the one hand, the fig mating system fosters reproductive isolation by promoting host specificity and limiting opportunities for heterospecific pollen transfer, but pollinator sharing nonetheless occurs and leads to hybridization and introgression between numerous fig species. On the other hand, the occurrence of host sharing and host switching creates opportunities for interspecific interactions among wasp species, yet hybridization and introgression is apparently rare or absent. Similarly, in the yucca and yucca moth system, another well-known obligate pollination mutualism, the host plants are known to share pollinators and to hybridize (Leebens-Mack et al., 1998; Smith et al., 2008; Rentsch and Leebens-Mack, 2012; Starr et al., 2013; Yoder et al., 2013; Arteaga et al., 2020), while the pollinators are genetically distinct and exhibit no evidence of hybridization (Leebens-Mack et al., 1998). As in the yucca pollinators, it appears hybridization and introgression of fig pollinator wasps have not been processes influencing their diversification or coevolution with their host plants.

Although hybridization has long been considered to be more prevalent in plants than in animals (Stebbins, 1959), there is a growing appreciation of the role hybridization has played in the animal tree of life (Mallet et al., 2016; Taylor and Larson, 2019). This is becoming more apparent as access to genomic data sets and newly developed statistical methods have improved our ability to detect signals of hybridization and introgression in the genomes of animal lineages. While we suspect a combination of pre- and postzygotic mechanisms limit successful hybridization and introgression in fig wasps, it is necessary to test these hypotheses with genome-scale data generated from communities of sympatric pollinator species. Observational experiments are also important in testing reproductive barriers between interacting species, however, successfully conducting such experiments with fig wasps requires overcoming significant challenges imposed by the closed structure of the syconium environment in which the heterospecific interactions of interest occur. Given the dearth of studies explicitly testing for hybridization and introgression in the pollinators—studies often focus on these processes in the host plants only—we hope future work will explicitly test for these processes in fig wasps. If the lack of hybridization and introgression demonstrated so far in fig wasps is the case in other fig systems, as we suspect, this will focus research on understanding the mechanisms promoting reproductive isolation in the pollinators in the face of frequent, and intimate, heterospecific interactions.

Machado et al. (2005) put forth the hypothesis that pollinator sharing and host switching have played prominent roles in generating the tremendous species diversity in the fig and wasp mutualism. In their model, genetically well-defined pollinator species move among genetically less well-defined fig species—either through pollinator sharing or host switching—and these opportunities for heterospecific pollination promote hybridization, introgression, and hybrid speciation between fig species. In this study, and others, F1 hybrid figs have been detected in nature. Because of the importance of fig floral volatiles for attracting host-specific pollinators (Van Noort et al., 1989; Ware et al., 1993; Grison-Pigé et al., 2002; Hossaert-McKey et al., 2010; Cornille et al., 2012), if the admixed volatile phenotypes produced by F1 hybrids attract pollinators (bearing compatible pollen) that are able to enter, pollinate, and reproduce inside the syconium, then generations of advanced generation hybrids may be formed, potentially providing a mechanism for adaptive interspecific gene flow and, possibly, hybrid speciation and diversification in the figs. And if pollinator wasps become consistently attracted to the volatile blends produced by the hybrid individuals, then this provides a mechanism for pollinator sharing or host switching. In the case of a host shift, reproductive isolation and diversification would be promoted in the wasps. Given the observations made within this Panamanian free-standing fig community, our results provide support for the Machado et al. (2005) model and suggest divergent evolutionary processes are responsible for generating diversification in the figs and their pollinator wasps.

## Conclusions

Consistent with findings in other sections of *Ficus*, we demonstrate hybridization at shallow time-scales between multiple fig species and introgression at deep time-scales between two of the five *Pharmacosycea* fig species. In contrast, the six *Tetrapus* pollinating wasps in this community are genetically distinct, well-defined species, and show no evidence of hybridization or introgression, consistent with findings from other fig and fig wasp systems. Although this obligate mutualism is maintained by tight ecological associations, processes affecting diversification differ between host and pollinator. Our findings are consistent with observations in other obligate pollination mutualisms, and suggest hybridization and introgression are processes affecting the evolution of the host plants, but not of their associated pollinators.

## Acknowledgements

We thank Juan C Penagos Zuluaga, Brian Park, Finn Piatscheck, and Jose Lopez for extracting DNA from the host figs. We thank the Heath Lab for helpful comments and discussion that improved the manuscript. Computational resources were provided by ResearchIT and the College of Liberal Arts and Sciences at Iowa State University. The Smithsonian Tropical Research Institute provided the research facilities and intellectual community that helped make this study possible.

## Data Accessibility

Raw sequence data will be made available from the NCBI Sequence Read Archive (SRA) under BioProject ID: ### (###). All data sets and custom scripts will be made available on Dryad.

## Supplemental Information for

**Figure S1:**
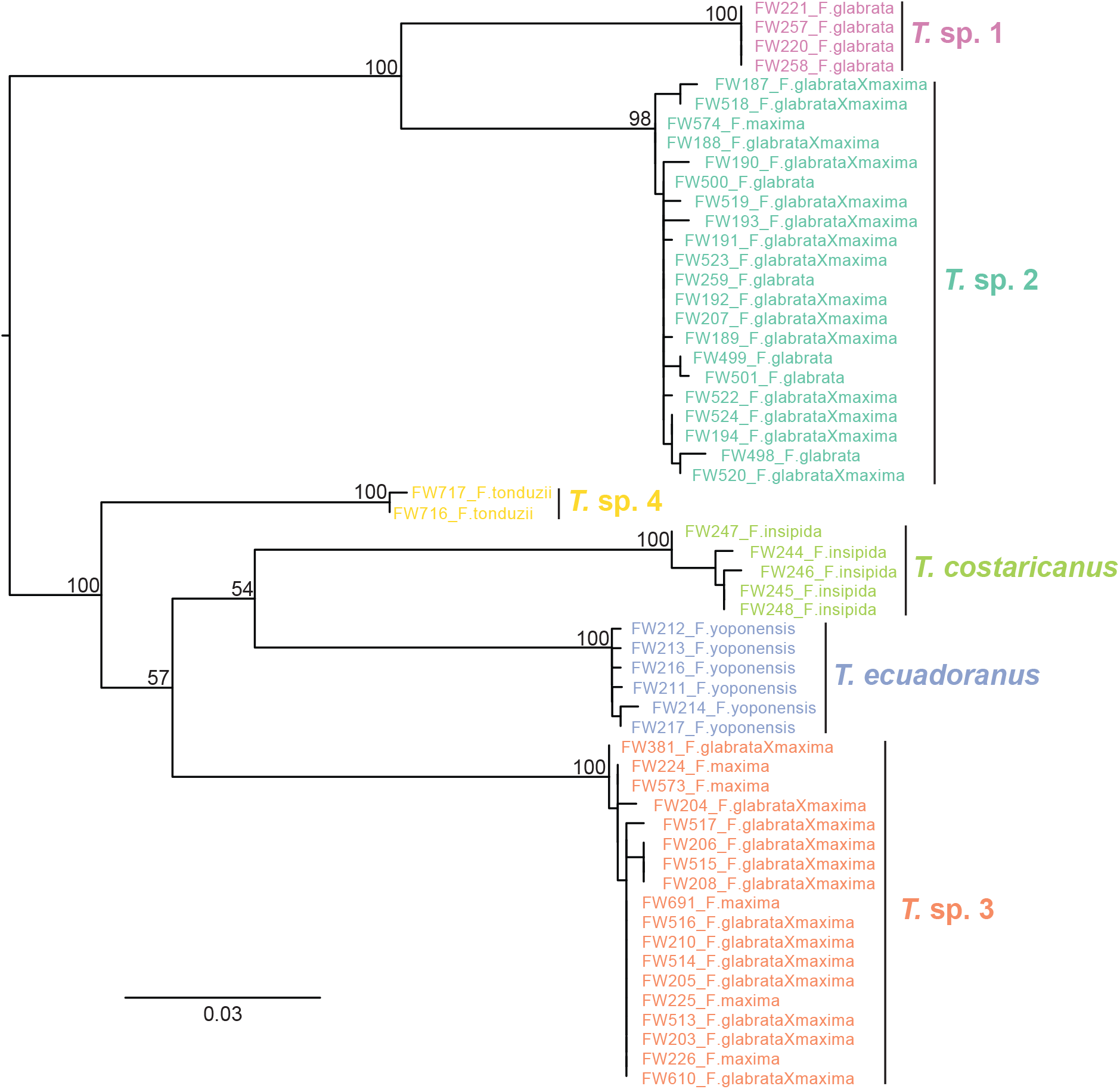
Maximum likelihood gene tree estimate of *Tetrapus* wasp COI data obtained from UCE sequencing. Nodal support values represent bootstrap support. Individuals are labeled with their unique identifier (FW#) and the host species from which they were sampled. The tree was midpoint rooted for visual purposes.

**Figure S2:**
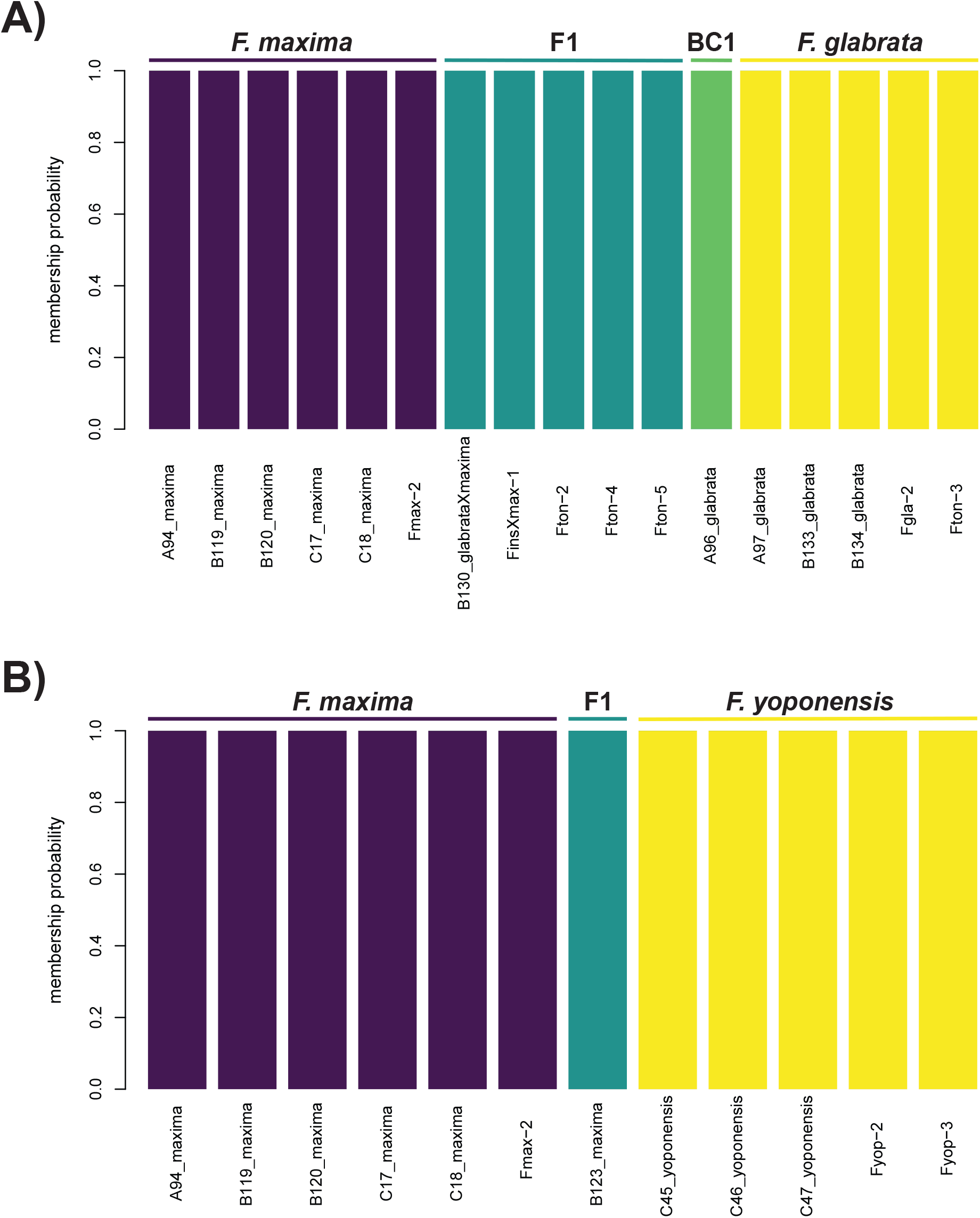
Hybrid analyses with snapclust. *F. glabrata* and *F. maxima*, including five F1s and one first-generation backcross (BC1) to *F. glabrata*. Panel B shows an F1 hybrid identified between *F. maxima* and *F. yoponensis*.

**Figure S3:**
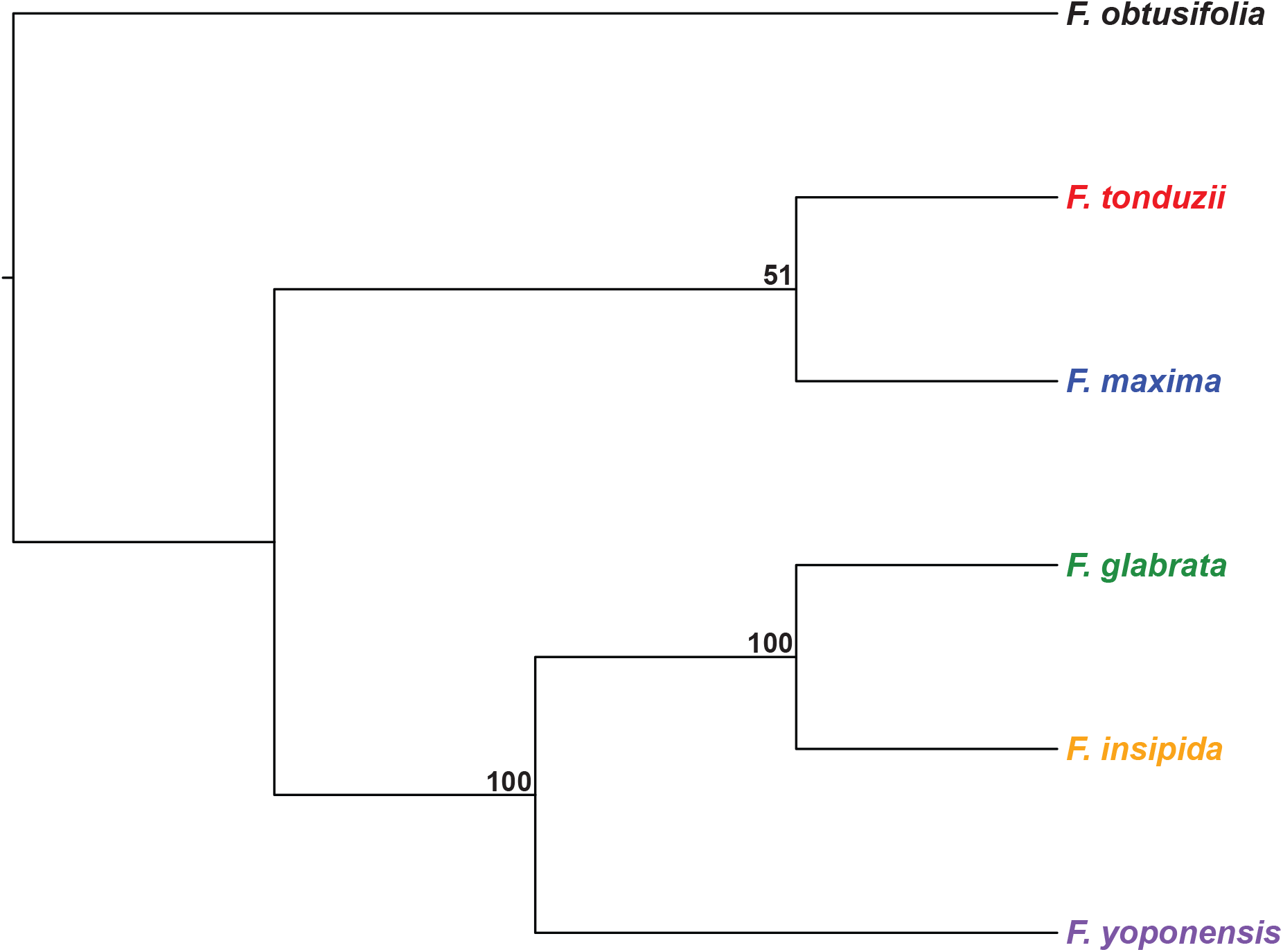
Species tree analysis with SVDQuartets of the *Pharmacosycea* figs. Nodal support values represent bootstrap support.

**Table S1:**
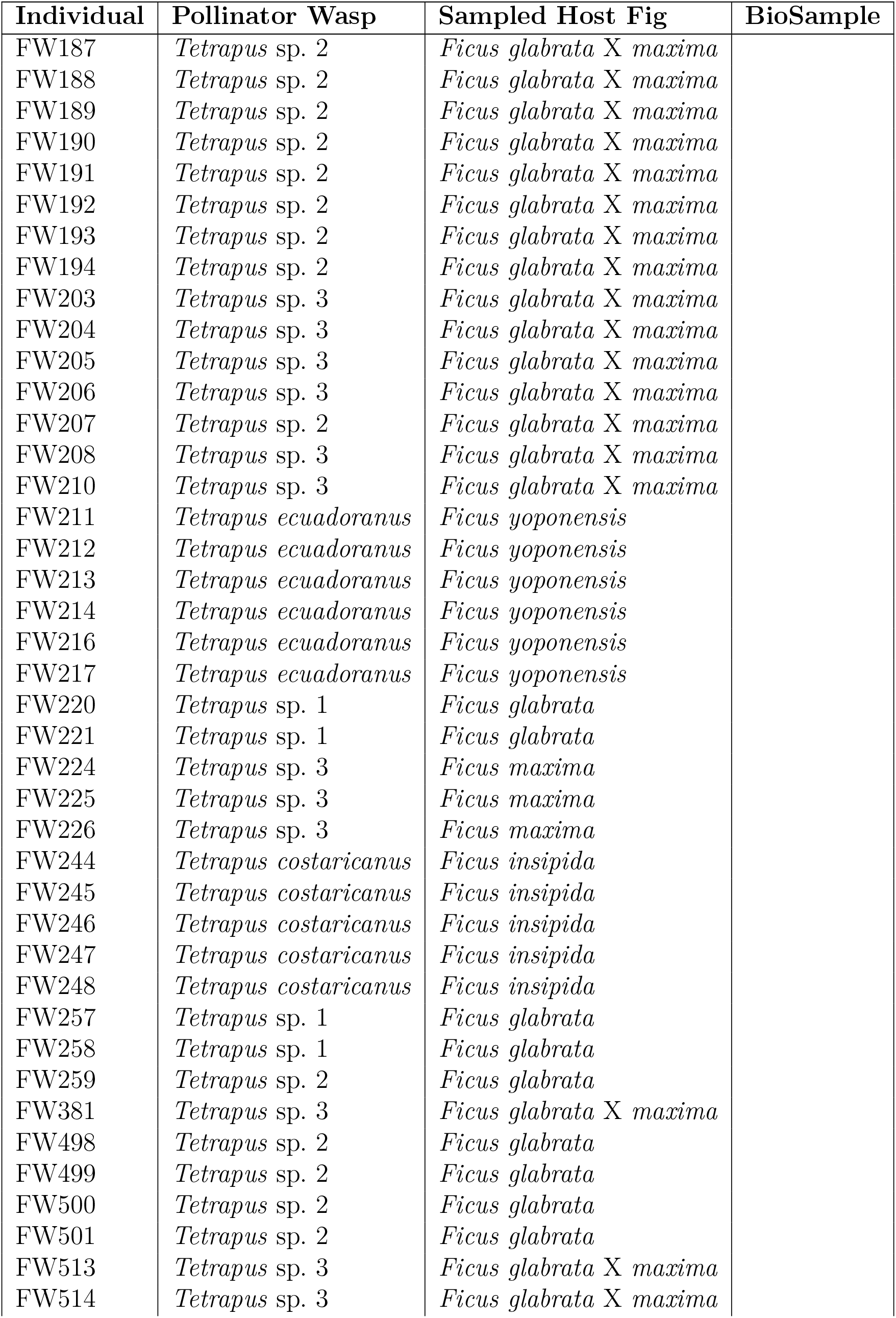
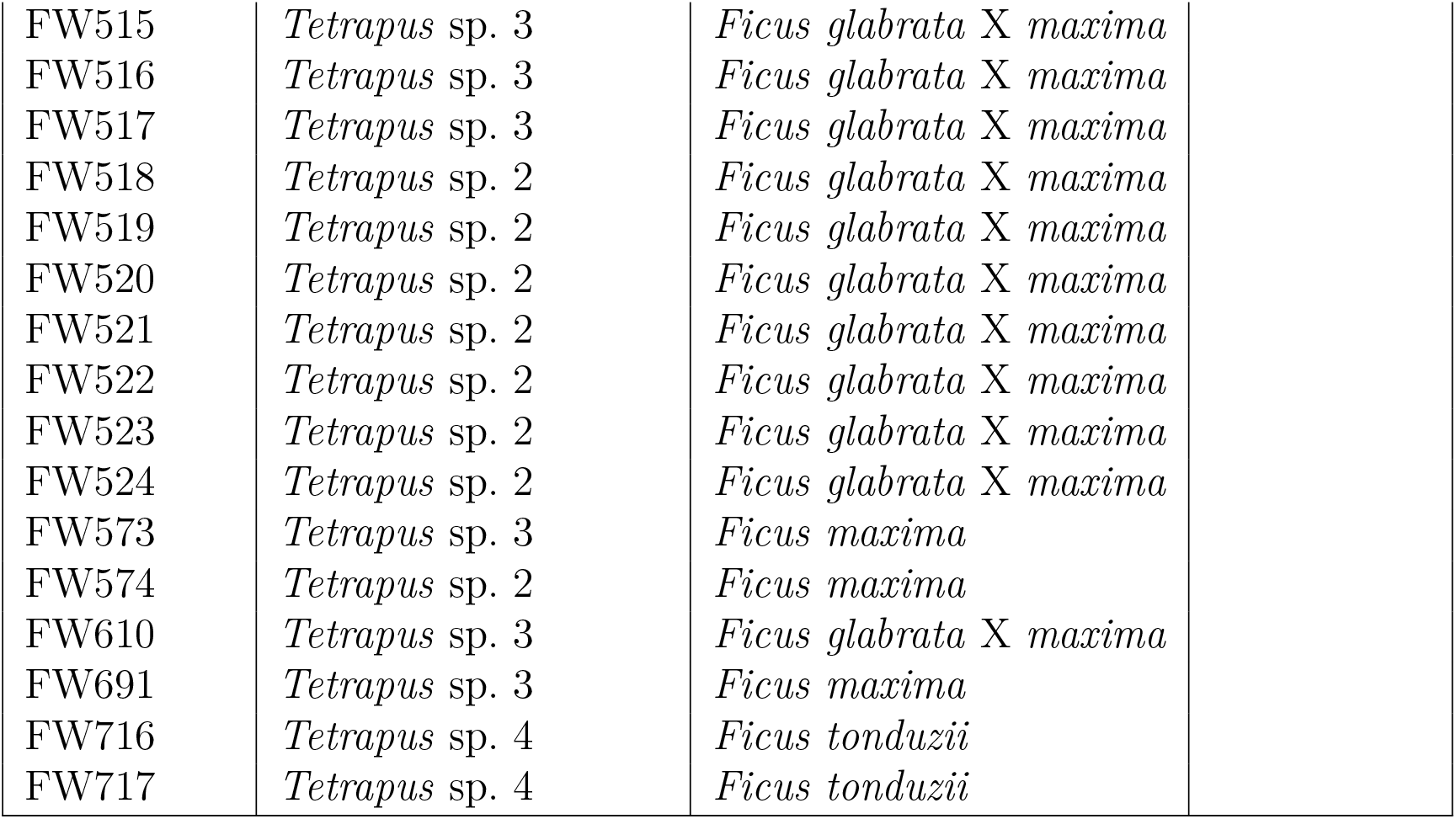
Pollinator wasp sampling. Information includes unique individual identifier (FW#), pollinator wasp species, sampled host fig species, and NCBI BioSample accession number.

| Individual | Pollinator Wasp | Sampled Host Fig | BioSample |
| --- | --- | --- | --- |
| FW187 | *Tetrapus* sp. 2 | *Ficus glabrata* X *maxima* |  |
| FW188 | *Tetrapus* sp. 2 | *Ficus glabrata* X *maxima* |  |
| FW189 | *Tetrapus* sp. 2 | *Ficus glabrata* X *maxima* |  |
| FW190 | *Tetrapus* sp. 2 | *Ficus glabrata* X *maxima* |  |
| FW191 | *Tetrapus* sp. 2 | *Ficus glabrata* X *maxima* |  |
| FW192 | *Tetrapus* sp. 2 | *Ficus glabrata* X *maxima* |  |
| FW193 | *Tetrapus* sp. 2 | *Ficus glabrata* X *maxima* |  |
| FW194 | *Tetrapus* sp. 2 | *Ficus glabrata* X *maxima* |  |
| FW203 | *Tetrapus* sp. 3 | *Ficus glabrata* X *maxima* |  |
| FW204 | *Tetrapus* sp. 3 | *Ficus glabrata* X *maxima* |  |
| FW205 | *Tetrapus* sp. 3 | *Ficus glabrata* X *maxima* |  |
| FW206 | *Tetrapus* sp. 3 | *Ficus glabrata* X *maxima* |  |
| FW207 | *Tetrapus* sp. 2 | *Ficus glabrata* X *maxima* |  |
| FW208 | *Tetrapus* sp. 3 | *Ficus glabrata* X *maxima* |  |
| FW210 | *Tetrapus* sp. 3 | *Ficus glabrata* X *maxima* |  |
| FW211 | *Tetrapus ecuadoranus* | *Ficus yoponensis* |  |
| FW212 | *Tetrapus ecuadoranus* | *Ficus yoponensis* |  |
| FW213 | *Tetrapus ecuadoranus* | *Ficus yoponensis* |  |
| FW214 | *Tetrapus ecuadoranus* | *Ficus yoponensis* |  |
| FW216 | *Tetrapus ecuadoranus* | *Ficus yoponensis* |  |
| FW217 | *Tetrapus ecuadoranus* | *Ficus yoponensis* |  |
| FW220 | *Tetrapus* sp. 1 | *Ficus glabrata* |  |
| FW221 | *Tetrapus* sp. 1 | *Ficus glabrata* |  |
| FW224 | *Tetrapus* sp. 3 | *Ficus maxima* |  |
| FW225 | *Tetrapus* sp. 3 | *Ficus maxima* |  |
| FW226 | *Tetrapus* sp. 3 | *Ficus maxima* |  |
| FW244 | *Tetrapus costaricanus* | *Ficus insipida* |  |
| FW245 | *Tetrapus costaricanus* | *Ficus insipida* |  |
| FW246 | *Tetrapus costaricanus* | *Ficus insipida* |  |
| FW247 | *Tetrapus costaricanus* | *Ficus insipida* |  |
| FW248 | *Tetrapus costaricanus* | *Ficus insipida* |  |
| FW257 | *Tetrapus* sp. 1 | *Ficus glabrata* |  |
| FW258 | *Tetrapus* sp. 1 | *Ficus glabrata* |  |
| FW259 | *Tetrapus* sp. 2 | *Ficus glabrata* |  |
| FW381 | *Tetrapus* sp. 3 | *Ficus glabrata* X *maxima* |  |
| FW498 | *Tetrapus* sp. 2 | *Ficus glabrata* |  |
| FW499 | *Tetrapus* sp. 2 | *Ficus glabrata* |  |
| FW500 | *Tetrapus* sp. 2 | *Ficus glabrata* |  |
| FW501 | *Tetrapus* sp. 2 | *Ficus glabrata* |  |
| FW513 | *Tetrapus* sp. 3 | *Ficus glabrata* X *maxima* |  |
| FW514 | *Tetrapus* sp. 3 | *Ficus glabrata* X *maxima* |  |

|  |  |  |
| --- | --- | --- |
| FW515 | *Tetrapus* sp. 3 | *Ficus glabrata* X *maxima* |
| FW516 | *Tetrapus* sp. 3 | *Ficus glabrata* X *maxima* |
| FW517 | *Tetrapus* sp. 3 | *Ficus glabrata* X *maxima* |
| FW518 | *Tetrapus* sp. 2 | *Ficus glabrata* X *maxima* |
| FW519 | *Tetrapus* sp. 2 | *Ficus glabrata* X *maxima* |
| FW520 | *Tetrapus* sp. 2 | *Ficus glabrata* X *maxima* |
| FW521 | *Tetrapus* sp. 2 | *Ficus glabrata* X *maxima* |
| FW522 | *Tetrapus* sp. 2 | *Ficus glabrata* X *maxima* |
| FW523 | *Tetrapus* sp. 2 | *Ficus glabrata* X *maxima* |
| FW524 | *Tetrapus* sp. 2 | *Ficus glabrata* X *maxima* |
| FW573 | *Tetrapus* sp. 3 | *Ficus maxima* |
| FW574 | *Tetrapus* sp. 2 | *Ficus maxima* |
| FW610 | *Tetrapus* sp. 3 | *Ficus glabrata* X *maxima* |
| FW691 | *Tetrapus* sp. 3 | *Ficus maxima* |
| FW716 | *Tetrapus* sp. 4 | *Ficus tonduzii* |
| FW717 | *Tetrapus* sp. 4 | *Ficus tonduzii* |

**Table S2:** Host fig sampling. Information includes unique individual identifier (F#), host fig species, and NCBI BioSample accession number.

| Individual | Sampled Host Fig | BioSample |
| --- | --- | --- |
| F1 | *Ficus glabrata* X *maxima* |  |
| F2 | *Ficus glabrata* |  |
| F3 | *Ficus glabrata* |  |
| F4 | *Ficus glabrata* |  |
| F5 | *Ficus glabrata* |  |
| F6 | *Ficus glabrata* X *maxima* |  |
| F7 | *Ficus insipida* |  |
| F8 | *Ficus insipida* |  |
| F9 | *Ficus insipida* |  |
| F10 | *Ficus insipida* |  |
| F11 | *Ficus insipida* |  |
| F12 | *Ficus insipida* |  |
| F13 | *Ficus glabrata* X *maxima* |  |
| F14 | *Ficus maxima* |  |
| F15 | *Ficus maxima* |  |
| F16 | *Ficus maxima* |  |
| F17 | *Ficus maxima* X *yoponensis* |  |
| F18 | *Ficus maxima* |  |
| F19 | *Ficus maxima* |  |
| F20 | *Ficus maxima* |  |
| F21 | *Ficus tonduzii* |  |
| F22 | *Ficus glabrata* X *maxima* |  |
| F23 | *Ficus glabrata* |  |
| F24 | *Ficus glabrata* X *maxima* |  |
| F25 | *Ficus glabrata* X *maxima* |  |
| F26 | *Ficus yoponensis* |  |
| F27 | *Ficus yoponensis* |  |
| F28 | *Ficus yoponensis* |  |
| F29 | *Ficus yoponensis* |  |
| F30 | *Ficus yoponensis* |  |

**Table S3:**
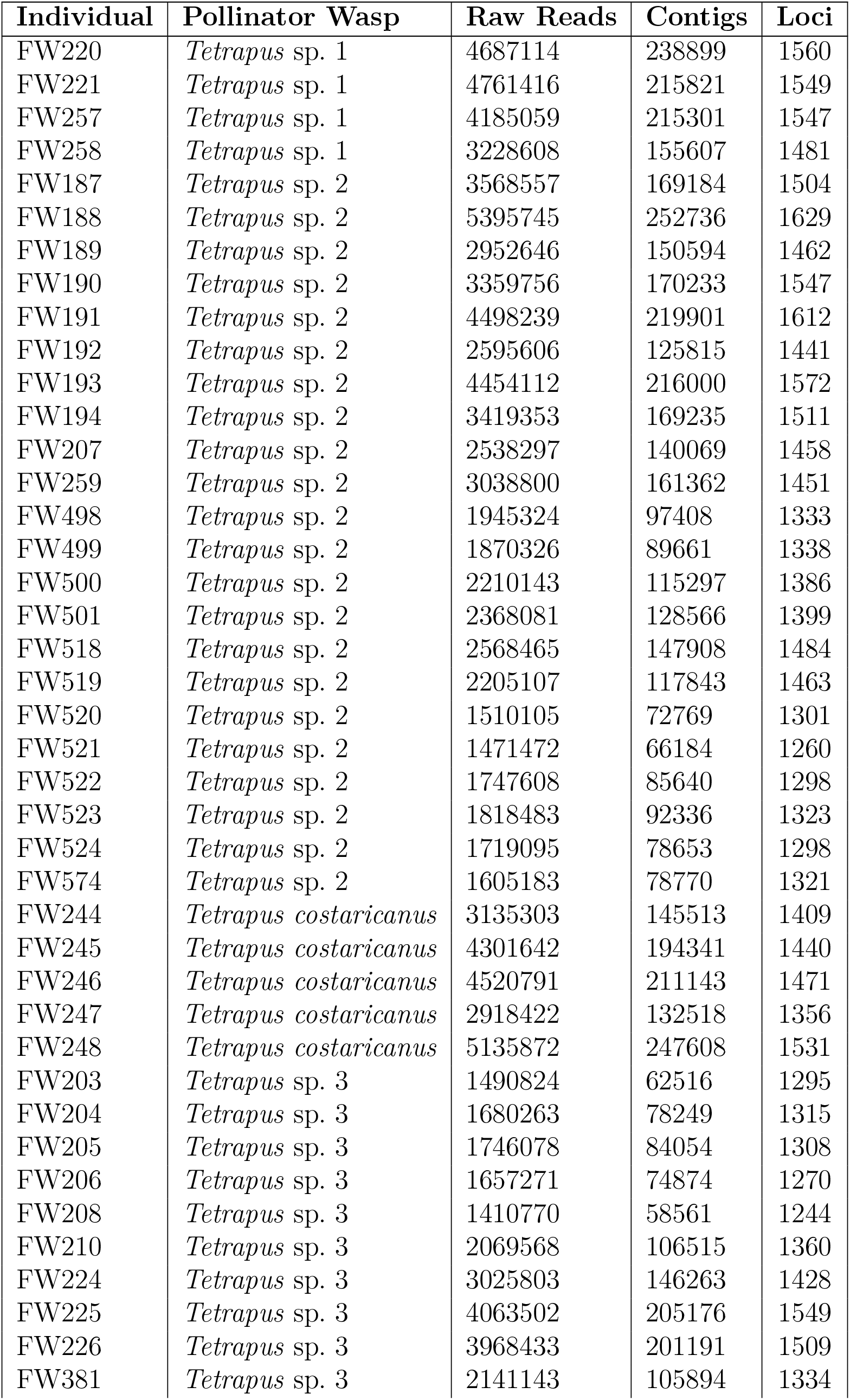
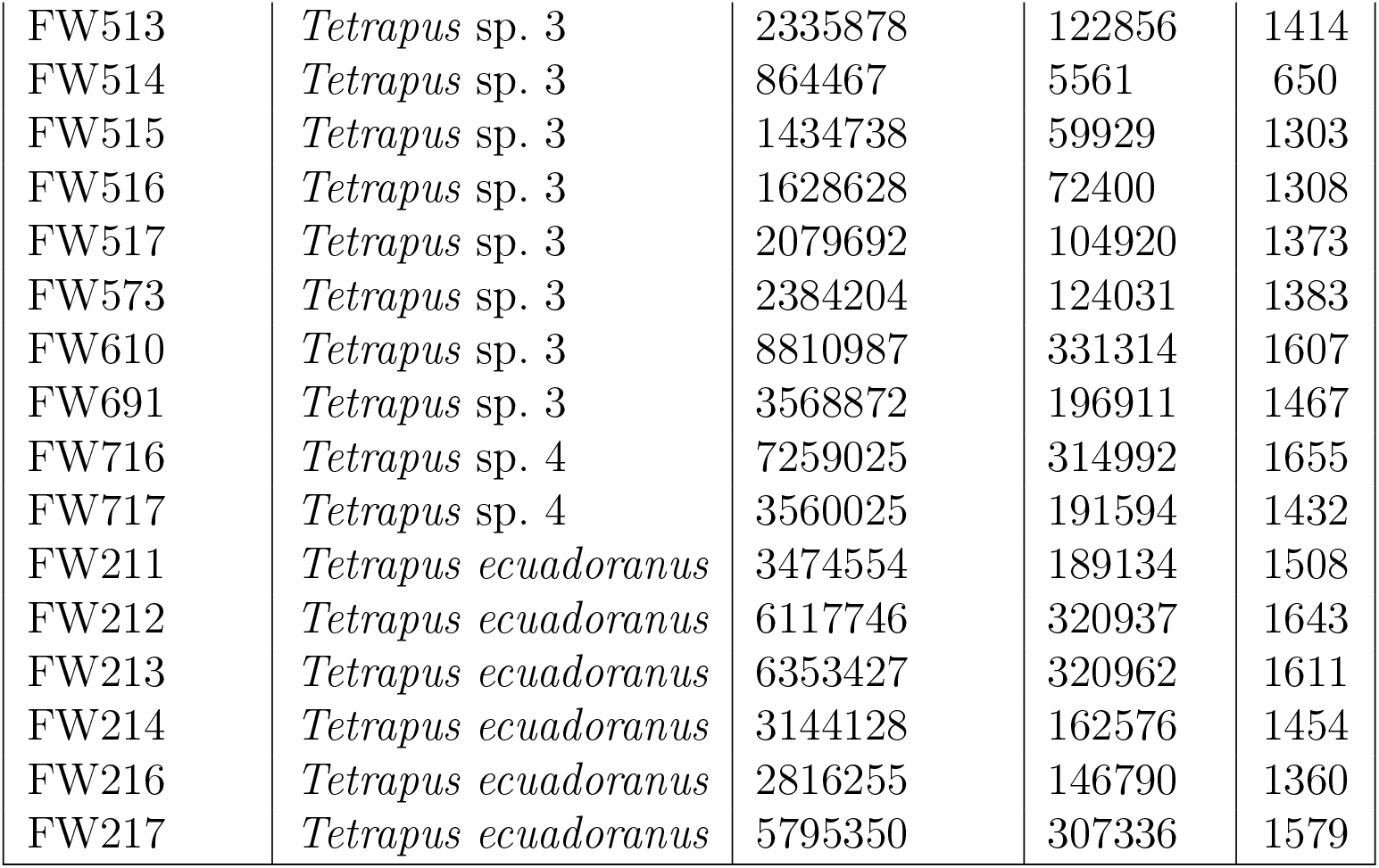
Pollinator wasp sequencing. Information includes unique individual identifier (FW#), pollinator wasp species, number of raw sequence reads, number of Trinity contigs, and number of UCE loci.

**Table S4:** Host fig sequencing. Information includes unique individual identifier (F#), host fig species, number of raw sequence reads, number of RAD clusters, and number of RAD loci when requiring at least 50% sampling presence.

| Individual | Host Fig | Raw Reads | Clusters | Loci |
| --- | --- | --- | --- | --- |
| F1 | *Ficus glabrata* X *maxima* | 7244559 | 176519 | 32601 |
| F2 | *Ficus glabrata* | 4887963 | 165416 | 24792 |
| F3 | *Ficus glabrata* | 4214450 | 63549 | 34655 |
| F4 | *Ficus glabrata* | 7119332 | 72796 | 36369 |
| F5 | *Ficus glabrata* | 2583903 | 60137 | 20939 |
| F6 | *Ficus glabrata* X *maxima* | 4060762 | 67775 | 35566 |
| F7 | *Ficus insipida* | 1003795 | 44201 | 16507 |
| F8 | *Ficus insipida* | 7909847 | 76907 | 36970 |
| F9 | *Ficus insipida* | 2017468 | 50610 | 29427 |
| F10 | *Ficus insipida* | 6030721 | 74683 | 36611 |
| F11 | *Ficus insipida* | 6928368 | 67783 | 36858 |
| F12 | *Ficus insipida* | 7347462 | 77847 | 37166 |
| F13 | *Ficus glabrata* X *maxima* | 1772506 | 36250 | 16900 |
| F14 | *Ficus maxima* | 2082861 | 53092 | 29079 |
| F15 | *Ficus maxima* | 2054747 | 63114 | 27703 |
| F16 | *Ficus maxima* | 2311959 | 57340 | 31056 |
| F17 | *Ficus maxima* X *yoponensis* | 1547830 | 47958 | 23657 |
| F18 | *Ficus maxima* | 24089307 | 93134 | 37423 |
| F19 | *Ficus maxima* | 15711473 | 81113 | 37464 |
| F20 | *Ficus maxima* | 1917990 | 40238 | 17481 |
| F21 | *Ficus tonduzii* | 20840966 | 116802 | 36924 |
| F22 | *Ficus glabrata* X *maxima* | 15431214 | 86425 | 37364 |
| F23 | *Ficus glabrata* | 19882495 | 92835 | 37502 |
| F24 | *Ficus glabrata* X *maxima* | 17770360 | 95279 | 37868 |
| F25 | *Ficus glabrata* X *maxima* | 1947746 | 37936 | 18174 |
| F26 | *Ficus yoponensis* | 25289469 | 97790 | 37606 |
| F27 | *Ficus yoponensis* | 17247374 | 84773 | 37546 |
| F28 | *Ficus yoponensis* | 10196819 | 74354 | 37302 |
| F29 | *Ficus yoponensis* | 10046758 | 70781 | 35725 |
| F30 | *Ficus yoponensis* | 12440898 | 78933 | 36314 |

## References

Ahmed, S., S. G. Compton, R. K. Butlin, and P. M. Gilmartin. 2009. Wind-borne insects mediate directional pollen transfer between desert fig trees 160 kilometers apart. Proceedings of the National Academy of Sciences 106:20342–20347.

Andermann, T., A. M. Fernandes, U. Olsson, M. Töpel, B. Pfeil, B. Oxelman, A. Aleixo, B. C. Faircloth, and A. Antonelli. 2018. Allele phasing greatly improves the phylogenetic utility of ultraconserved elements. Systematic Biology 68:32–46.

Anderson, E. 1953. Introgressive hybridization. Biological reviews of the cambridge philosophical society 28:280–307.

Arteaga, M. C., R. Bello-Bedoy, and J. Gasca-Pineda. 2020. Hybridization between yuccas from Baja California: genomic and environmental patterns. Frontiers in Plant Science 11:685.

Baird, N. A., P. D. Etter, T. S. Atwood, M. C. Currey, A. L. Shiver, Z. A. Lewis, E. U. Selker, W. A. Cresko, and E. A. Johnson. 2008. Rapid SNP discovery and genetic mapping using sequenced RAD markers. PLoS ONE 3:e3376.

Berg, C. 1989. Classification and distribution of Ficus. Experientia 45:605–611.

Berg, C. 2006. The subdivision of Ficus subgenus Pharmacosycea section Pharmacosycea (Moraceae). Blumea 51:147–151.

Beugin, M.-P., T. Gayet, D. Pontier, S. Devillard, and T. Jombart. 2018. A fast likelihood solution to the genetic clustering problem. Methods in Ecology and Evolution 9:1006–1016.

Bolger, A. M., M. Lohse, and B. Usadel. 2014. Trimmomatic: a flexible trimmer for Illumina sequence data. Bioinformatics 30:2114–2120.

Branstetter, M. G., J. T. Longino, P. S. Ward, and B. C. Faircloth. 2017. Enriching the ant tree of life: enhanced uce bait set for genome-scale phylogenetics of ants and other hymenoptera. Methods in Ecology and Evolution 8:768–776.

Bronstein, J. L. 1987. Maintenance of species-specificity in a neotropical fig–pollinator wasp mutualism. Oikos 48:39–46.

Bruun-Lund, S., W. L. Clement, F. Kjellberg, and N. Rønsted. 2017. First plastid phylogenomic study reveals potential cyto-nuclear discordance in the evolutionary history of Ficus L.(Moraceae). Molecular Phylogenetics and Evolution 109:93–104.

Castresana, J. 2000. Selection of conserved blocks from multiple alignments for their use in phylogenetic analysis. Molecular Biology and Evolution 17:540–552.

Chernomor, O., A. Von Haeseler, and B. Q. Minh. 2016. Terrace aware data structure for phylogenomic inference from supermatrices. Systematic Biology 65:997–1008.

Chifman, J., and L. Kubatko. 2014. Quartet inference from SNP data under the coalescent model. Bioinformatics 30:3317–3324.

Compton, S. G. 1990. A collapse of host specificity in some African fig wasps. South African Journal of Science 86:39–40.

Cook, J. M., and J.-Y. Rasplus. 2003. Mutualists with attitude: coevolving fig wasps and figs. Trends in Ecology & Evolution 18:241–248.

Cornille, A., J. Underhill, A. Cruaud, M. Hossaert-McKey, S. Johnson, K. Tolley, F. Kjellberg, S. Van Noort, and M. Proffit. 2012. Floral volatiles, pollinator sharing and diversification in the fig–wasp mutualism: insights from Ficus natalensis, and its two wasp pollinators (South Africa). Proceedings of the Royal Society of London B: Biological Sciences 279:1731–1739.

Croat, T. B. 1978. Flora of Barro Colorado Island. Stanford University Press.

Cruaud, A., N. RÃÿnsted, B. Chantarasuwan, L. S. Chou, W. L. Clement, A. Couloux, B. Cousins, G. Genson, R. D. Harrison, P. E. Hanson, M. Hossaert-Mckey, R. Jabbour-Zahab, E. Jousselin, C. KerdelhuÃr, F. Kjellberg, C. Lopez-Vaamonde, J. Peebles, Y.-Q. Peng, R. A. S. Pereira, T. Schramm, R. Ubaidillah, S. van Noort, G. D. Weiblen, D.-R. Yang, A. Yodpinyanee, R. Libeskind-Hadas, J. M. Cook, J.-Y. Rasplus, and V. Savolainen. 2012. An extreme case of plant–insect codiversification: figs and fig-pollinating wasps. Systematic Biology 61:1029–1047.

Darwell, C. T., S. al Beidh, and J. M. Cook. 2014. Molecular species delimitation of a symbiotic fig-pollinating wasp species complex reveals extreme deviation from reciprocal partner specificity. BMC Evolutionary Biology 14:189.

de Medeiros, B. A., and B. D. Farrell. 2020. Evaluating insect-host interactions as a driver of species divergence in palm flower weevils. Communications Biology 3:1–9.

Dierckxsens, N., P. Mardulyn, and G. Smits. 2017. Novoplasty: de novo assembly of organelle genomes from whole genome data. Nucleic Acids Research 45:e18.

Doyle, J. J., and J. L. Doyle. 1987. A rapid DNA isolation procedure for small quantities of fresh leaf tissue. Phytochem Bull 19:11–15.

Eaton, D. A., A. L. Hipp, A. González-Rodríguez, and J. Cavender-Bares. 2015. Historical introgression among the American live oaks and the comparative nature of tests for introgression. Evolution 69:2587–2601.

Eaton, D. A., and I. Overcast. 2020. ipyrad: Interactive assembly and analysis of RADseq datasets. Bioinformatics 36:2592–2594.

Edelman, N. B., P. B. Frandsen, M. Miyagi, B. Clavijo, J. Davey, R. B. Dikow, G. GarcÃ ŋa-Accinelli, S. M. V. Belleghem, N. Patterson, D. E. Neafsey, R. Challis, S. Kumar, G. R. P. Moreira, C. Salazar, M. Chouteau, B. A. Counterman, R. Papa, M. Blaxter, R. D. Reed, K. K. Dasmahapatra, M. Kronforst, M. Joron, C. D. Jiggins, W. O. McMillan, F. D. Palma, A. J. Blumberg, J. Wakeley, D. Jaffe, and J. Mallet. 2019. Genomic architecture and introgression shape a butterfly radiation. Science 366:594–599.

Edelman, N. B., and J. Mallet. 2021. Prevalence and adaptive impact of introgression. Annual Review of Genetics 55:265–283.

Enciso-Romero, J., C. Pardo-Díaz, S. H. Martin, C. F. Arias, M. Linares, W. O. McMillan, C. D. Jiggins, and C. Salazar. 2017. Evolution of novel mimicry rings facilitated by adaptive introgression in tropical butterflies. Molecular Ecology 26:5160–5172.

Faircloth, B. C. 2013. Illumiprocessor: a trimmomatic wrapper for parallel adapter and quality trimming. See http://dx.doi.org/10.6079/J9ILL (accessed 4 November 2016).

Faircloth, B. C. 2015. PHYLUCE is a software package for the analysis of conserved genomic loci. Bioinformatics 32:786–788.

Francis, R. M. 2017. pophelper: an r package and web app to analyse and visualize population structure. Molecular Ecology Resources 17:27–32.

Glenn, T. C., R. A. Nilsen, T. J. Kieran, J. G. Sanders, N. J. Bayona-Vásquez, J. W. Finger, T. W. Pierson, K. E. Bentley, S. L. Hoffberg, S. Louha, F. J. Garcia-De Leon, M. A. del Rio Portilla, K. D. Reed, J. L. Anderson, J. K. Meece, S. E. Aggrey, R. Rekaya, M. Alabady, M. Belanger, K. Winker, and B. C. Faircloth. 2019. Adapterama i: universal stubs and primers for 384 unique dual-indexed or 147,456 combinatorially-indexed illumina libraries (itru & inext). PeerJ 7:e7755.

Grabherr, M. G., B. J. Haas, M. Yassour, J. Z. Levin, D. A. Thompson, I. Amit, X. Adiconis, L. Fan, R. Raychowdhury, Q. Zeng, Z. Chen, E. Mauceli, N. Hacohen, A. Gnirke, N. Rhind, F. di Palma, B. W. Birren, C. Nusbaum, K. Lindblad-Toh, N. Friedman, and A. Regev. 2011. Full-length transcriptome assembly from RNA-Seq data without a reference genome. Nature Biotechnology 29:644–652.

Grison-Pigé, L., J.-M. Bessière, and M. Hossaert-McKey. 2002. Specific attraction of figpollinating wasps: role of volatile compounds released by tropical figs. Journal of Chemical Ecology 28:283–295.

Hamilton, J. A., and J. M. Miller. 2016. Adaptive introgression as a resource for management and genetic conservation in a changing climate. Conservation Biology 30:33–41.

Hedrick, P. W. 2013. Adaptive introgression in animals: examples and comparison to new mutation and standing variation as sources of adaptive variation. Molecular Ecology 22:4606– 4618.

Hembry, D. H., and D. M. Althoff. 2016. Diversification and coevolution in brood pollination mutualisms: windows into the role of biotic interactions in generating biological diversity. American Journal of Botany 103:1783–1792.

Hembry, D. H., A. Kawakita, N. E. Gurr, M. A. Schmaedick, B. G. Baldwin, and R. G. Gillespie. 2013. Non-congruent colonizations and diversification in a coevolving pollination mutualism on oceanic islands. Proceedings of the Royal Society B: Biological Sciences 280:20130361.

Herre, E. A. 1989. Coevolution of reproductive characteristics in 12 species of New World figs and their pollinator wasps. Experientia 45:637–647.

Herre, E. A., K. C. Jandér, and C. A. Machado. 2008. Evolutionary ecology of figs and their associates: recent progress and outstanding puzzles. Annual Review of Ecology, Evolution, and Systematics 39:439–458.

Hoang, D. T., O. Chernomor, A. Von Haeseler, B. Q. Minh, and L. S. Vinh. 2018. Ufboot2: improving the ultrafast bootstrap approximation. Molecular Biology and Evolution 35:518– 522.

Hossaert-McKey, M., C. Soler, B. Schatz, and M. Proffit. 2010. Floral scents: their roles in nursery pollination mutualisms. Chemoecology 20:75–88.

Jackson, A. P., C. A. Machado, N. Robbins, and E. A. Herre. 2008. Multi-locus phylogenetic analysis of neotropical figs does not support co-speciation with the pollinators: the importance of systematic scale in fig/wasp cophylogenetic studies. Symbiosis 45:57–72.

Janzen, D. H. 1979. How to be a fig. Annual Review of Ecology and Systematics 10:13–51.

Jombart, T. 2008. adegenet: a r package for the multivariate analysis of genetic markers. Bioinformatics 24:1403–1405.

Kalyaanamoorthy, S., B. Q. Minh, T. K. Wong, A. von Haeseler, and L. S. Jermiin. 2017. Modelfinder: fast model selection for accurate phylogenetic estimates. Nature Methods 14:587–589.

Katoh, K., and D. M. Standley. 2013. MAFFT multiple sequence alignment software version 7: improvements in performance and usability. Molecular Biology and Evolution 30:772– 780.

Leebens-Mack, J., O. Pellmyr, and M. Brock. 1998. Host specificity and the genetic structure of two Yucca moth species in a Yucca hybrid zone. Evolution 52:1376–1382.

Leroy, T., J.-M. Louvet, C. Lalanne, G. Le Provost, K. Labadie, J.-M. Aury, S. Delzon, C. Plomion, and A. Kremer. 2020. Adaptive introgression as a driver of local adaptation to climate in European white oaks. New Phytologist 226:1171–1182.

Li, H., and R. Durbin. 2010. Fast and accurate short read alignment with Burrows–Wheeler transform. Bioinformatics 26:589–595.

Li, H., B. Handsaker, A. Wysoker, T. Fennell, J. Ruan, N. Homer, G. Marth, G. Abecasis, and R. Durbin. 2009. The sequence alignment/map format and SAMtools. Bioinformatics 25:2078–2079.

Machado, C. A., E. Jousselin, F. Kjellberg, S. G. Compton, and E. A. Herre. 2001. Phylogenetic relationships, historical biogeography and character evolution of fig-pollinating wasps. Proceedings of the Royal Society of London B: Biological Sciences 268:685–694.

Machado, C. A., N. Robbins, M. T. P. Gilbert, and E. A. Herre. 2005. Critical review of host specificity and its coevolutionary implications in the fig/fig-wasp mutualism. Proceedings of the National Academy of Sciences 102:6558–6565.

Malinsky, M., H. Svardal, A. M. Tyers, E. A. Miska, M. J. Genner, G. F. Turner, and R. Durbin. 2018. Whole-genome sequences of Malawi cichlids reveal multiple radiations interconnected by gene flow. Nature Ecology & Evolution 2:1940–1955.

Mallet, J. 2005. Hybridization as an invasion of the genome. Trends in Ecology & Evolution 20:229–237.

Mallet, J., N. Besansky, and M. W. Hahn. 2016. How reticulated are species? BioEssays 38:140–149.

Marussich, W. A., and C. A. Machado. 2007. Host-specificity and coevolution among pollinating and nonpollinating new world fig wasps. Molecular Ecology 16:1925–1946.

McLeish, M. J., and S. Van Noort. 2012. Codivergence and multiple host species use by fig wasp populations of the Ficus pollination mutualism. BMC Evolutionary Biology 12.

McVay, J. D., A. L. Hipp, and P. S. Manos. 2017. A genetic legacy of introgression confounds phylogeny and biogeography in oaks. Proceedings of the Royal Society B: Biological Sciences 284:20170300.

Meier, J. I., D. A. Marques, S. Mwaiko, C. E. Wagner, L. Excoffier, and O. Seehausen. 2017. Ancient hybridization fuels rapid cichlid fish adaptive radiations. Nature Communications 8:1–11.

Moe, A. M., D. R. Rossi, and G. D. Weiblen. 2011. Pollinator sharing in dioecious figs (Ficus: Moraceae). Biological Journal of the Linnean Society 103:546–558.

Moe, A. M., and G. D. Weiblen. 2012. Pollinator-mediated reproductive isolation among dioecious fig species (Ficus, Moraceae). Evolution 66:3710–3721.

Molbo, D., C. A. Machado, E. A. Herre, and L. Keller. 2004. Inbreeding and population structure in two pairs of cryptic fig wasp species. Molecular Ecology 13:1613–1623.

Molbo, D., C. A. Machado, J. G. Sevenster, L. Keller, and E. A. Herre. 2003. Cryptic species of fig-pollinating wasps: implications for the evolution of the fig–wasp mutualism, sex allocation, and precision of adaptation. Proceedings of the National Academy of Sciences 100:5867–5872.

Nason, J. D., E. A. Herre, and J. L. Hamrick. 1996. Paternity analysis of the breeding structure of strangler fig populations: evidence for substantial long-distance wasp dispersal. Journal of Biogeography 23:501–512.

Nason, J. D., E. A. Herre, and J. L. Hamrick. 1998. The breeding structure of a tropical keystone plant resource. Nature 391:685–687.

Nguyen, L.-T., H. A. Schmidt, A. Von Haeseler, and B. Q. Minh. 2015. IQ-TREE: a fast and effective stochastic algorithm for estimating maximum-likelihood phylogenies. Molecular Biology and Evolution 32:268–274.

Olave, M., and A. Meyer. 2020. Implementing large genomic single nucleotide polymorphism data sets in phylogenetic network reconstructions: a case study of particularly rapid radiations of cichlid fish. Systematic Biology 69:848–862.

Pardo-Diaz, C., C. Salazar, S. W. Baxter, C. Merot, W. Figueiredo-Ready, M. Joron, W. O. McMillan, and C. D. Jiggins. 2012. Adaptive introgression across species boundaries in Heliconius butterflies. PLoS Genetics 8:e1002752.

Pellmyr, O., F. Kjellberg, E. A. Herre, A. Kawakita, D. H. Hembry, J. N. Holland, T. Terrazas, W. Clement, K. A. Segraves, and D. M. Althoff. 2020. Active pollination drives selection for reduced pollen-ovule ratios. American Journal of Botany 107:164–170.

Pickrell, J., and J. Pritchard. 2012. Inference of population splits and mixtures from genome-wide allele frequency data. PLoS Genetics 8:e1002967.

Pritchard, J. K., M. Stephens, and P. Donnelly. 2000. Inference of population structure using multilocus genotype data. Genetics 155:945–959.

R Core Team. 2018. R: A Language and Environment for Statistical Computing. R Foundation for Statistical Computing, Vienna, Austria. URL https://www.R-project.org/.

Raj, A., M. Stephens, and J. K. Pritchard. 2014. fastSTRUCTURE: variational inference of population structure in large SNP data sets. Genetics 197:573–589.

Ramírez, W. 1970. Host specificity of fig wasps (Agaonidae). Evolution 24:680–691.

Ramírez, W. 1994. Hybridization of Ficus religiosa with F. septica and F. aurea (Moraceae). Revista de biología tropical 42:339–342.

Renoult, J. P., F. Kjellberg, C. Grout, S. Santoni, and B. Khadari. 2009. Cyto-nuclear discordance in the phylogeny of Ficus section Galoglychia and host shifts in plant-pollinator associations. BMC Evolutionary Biology 9:248.

Rentsch, J. D., and J. Leebens-Mack. 2012. Homoploid hybrid origin of Yucca gloriosa: intersectional hybrid speciation in Yucca (agavoideae, asparagaceae). Ecology and Evolution 2:2213–2222.

Satler, J. D., E. A. Herre, T. A. Heath, C. A. Machado, A. G. Zúñiga, and J. D. Nason. 2022. Genome-wide sequence data show no evidence of hybridization and introgression among pollinator wasps associated with a community of Panamanian strangler figs. Molecular Ecology accepted.

Satler, J. D., E. A. Herre, K. C. Jandér, D. A. R. Eaton, C. A. Machado, T. A. Heath, and J. D. Nason. 2019. Inferring processes of coevolutionary diversification in a community of Panamanian strangler figs and associated pollinating wasps. Evolution 73:2295–2311.

Shanahan, M., S. So, S. G. Compton, and R. Corlett. 2001. Fig-eating by vertebrate frugivores: a global review. Biological Reviews 76:529–572.

Smith, C. I., W. K. Godsoe, S. Tank, J. B. Yoder, and O. Pellmyr. 2008. Distinguishing coevolution from covicariance in an obligate pollination mutualism: asynchronous divergence in Joshua tree and its pollinators. Evolution 62:2676–2687.

Solís-Lemus, C., and C. Ané. 2016. Inferring phylogenetic networks with maximum pseudolikelihood under incomplete lineage sorting. PLoS Genetics 12:e1005896.

Solís-Lemus, C., P. Bastide, and C. Ané. 2017. PhyloNetworks: a package for phylogenetic networks. Molecular Biology and Evolution 34:3292–3298.

Souto-Vilarós, D., A. Machac, J. Michalek, C. T. Darwell, M. Sisol, T. Kuyaiva, B. Isua, G. D. Weiblen, V. Novotny, and S. T. Segar. 2019. Faster speciation of fig-wasps than their host figs leads to decoupled speciation dynamics: snapshots across the speciation continuum. Molecular Ecology 28:3958–3976.

Starr, T. N., K. E. Gadek, J. B. Yoder, R. Flatz, and C. I. Smith. 2013. Asymmetric hybridization and gene flow between Joshua trees (Agavaceae: Yucca) reflect differences in pollinator host specificity. Molecular Ecology 22:437–449.

Stebbins, G. L. 1959. The role of hybridization in evolution. Proceedings of the American Philosophical Society 103:231–251.

Sutton, T. L., J. L. DeGabriel, M. Riegler, and J. M. Cook. 2017. Local coexistence and genetic isolation of three pollinator species on the same fig tree species. Heredity 118:486– 490.

Svardal, H., F. X. Quah, M. Malinsky, B. P. Ngatunga, E. A. Miska, W. Salzburger, M. J. Genner, G. F. Turner, and R. Durbin. 2020. Ancestral hybridization facilitated species diversification in the Lake Malawi cichlid fish adaptive radiation. Molecular Biology and Evolution 37:1100–1113.

Swofford, D. L. 2003. PAUP*: phylogenetic analysis using parsimony (*and other methods). Sinauer Associates, Sunderland, Massachusetts.

Taylor, S. A., and E. L. Larson. 2019. Insights from genomes into the evolutionary importance and prevalence of hybridization in nature. Nature Ecology & Evolution 3:170–177.

Van Noort, S., A. Ware, and S. Compton. 1989. Pollinator-specific volatile attractants released from the figs of Ficus burtt-davyi. South African Journal of Science 85:323–324.

Wang, G., C. H. Cannon, and J. Chen. 2016. Pollinator sharing and gene flow among closely related sympatric dioecious fig taxa. Proceedings of the Royal Society of London B: Biological Sciences 283:20152963.

Wang, G., X. Zhang, E. A. Herre, D. McKey, C. A. Machado, W.-B. Yu, C. H. Cannon, M. L. Arnold, R. A. Pereira, R. Ming, Y.-F. Liu, Y. Wang, D. Ma, and J. Chen. 2021a. Genomic evidence of prevalent hybridization throughout the evolutionary history of the fig-wasp pollination mutualism. Nature Communications 12:1–14.

Wang, R., Y. Yang, Y. Jing, S. T. Segar, Y. Zhang, G. Wang, J. Chen, Q.-F. Liu, S. Chen, Y. Chen, A. Cruaud, Y.-Y. Ding, D. W. Dunn, Q. Gao, P. M. Gilmartin, K. Jiang, F. Kjellberg, H.-Q. Li, Y.-Y. Li, J.-Q. Liu, M. Liu, C. A. Machado, R. Ming, J.-Y. Rasplus, X. Tong, P. Wen, H.-M. Yang, J.-J. Yang, Y. Yin, X.-T. Zhang, Y.-Y. Zhang, H. Yu, Z. Yue, S. G. Compton, and X.-Y. Chen. 2021b. Molecular mechanisms of mutualistic and antagonistic interactions in a plant–pollinator association. Nature Ecology & Evolution 5:974–986.

Ware, A. B., P. T. Kaye, S. G. Compton, and S. Van Noort. 1993. Fig volatiles: their role in attracting pollinators and maintaining pollinator specificity. Plant Systematics and Evolution 186:147–156.

Weiblen, G. D. 2002. How to be a fig wasp. Annual Review of Entomology 47:299–330.

Wilde, B. C., S. Rutherford, M. van der Merwe, M. L. Murray, and M. Rossetto. 2020. First example of hybridisation between two Australian figs (Moraceae). Australian Systematic Botany 33:436–445.

Yang, L.-Y., C. A. Machado, X.-D. Dang, Y.-Q. Peng, D.-R. Yang, D.-Y. Zhang, and W.-J. Liao. 2015. The incidence and pattern of copollinator diversification in dioecious and monoecious figs. Evolution 69:294–304.

Yoder, J., C. Smith, D. Rowley, R. Flatz, W. Godsoe, C. Drummond, and O. Pellmyr. 2013. Effects of gene flow on phenotype matching between two varieties of Joshua tree (Yucca brevifolia; agavaceae) and their pollinators. Journal of Evolutionary Biology 26:1220–1233.

Yu, H., E. Tian, L. Zheng, X. Deng, Y. Cheng, L. Chen, W. Wu, W. Tanming, D. Zhang, S. G. Compton, and F. Kjellberg. 2019. Multiple parapatric pollinators have radiated across a continental fig tree displaying clinal genetic variation. Molecular Ecology.

